# Cell Dynamic Mechanics Regulates Large-spatial Isotropic Matrix Modeling with Computational Simulations

**DOI:** 10.1101/2022.10.29.514382

**Authors:** Mingxing Ouyang, Yanling Hu, Weihui Chen, Hui Li, Yingbo Ji, Linshuo Qiu, Linlin Zhu, Baohua Ji, Bing Bu, Linhong Deng

**Author notes:** Corresponding authors: Dr. Mingxing Ouyang, Professor at Institute of Biomedical Engineering and Health Sciences, School of Medical and Health Engineering, Changzhou University, 1 Gehu Rd, Wujin District, Changzhou, 213164 China, Dr. Linhong Deng, Fellow of AIBME, Cheung Kung Distinguished Professor, Founding Director Institute of Biomedical Engineering and Health Sciences, School of Medical and Health Engineering, Changzhou University, 1 Gehu Rd, Wujin District, Changzhou, Jiangsu Province 213164 China, Dr. Bing Bu, Lecturer at Institute of Biomedical Engineering and Health Sciences, School of Medical and Health Engineering, Changzhou University, 1 Gehu Rd, Wujin District, Changzhou, 213164 China. Shared first authorships (Mingxing Ouyang, Yanling Hu).

## Abstract

Tissues are often isotropic and heterogeneous organizations, which developmental processes are coordinated by cells and extracellular matrix modeling. Cells have the capability of modeling matrix in distance, however, the biophysical mechanism is largely unknown. We investigated underlying mechanism of large collagen I (COL) fibrillary modeling by cell mechanics with designed arrays of cell clusters. By incorporating dynamic contractions, Molecular Dynamics simulations yielded highly matching isotropic outcomes with observed COL clustering in experiments from variable geometrical arrays without spatial limitation. Further designed single polygons from triangles to hexagons resulted in predicted structural assembly which showed maintained spatial balance. Cell cytoskeletal integrity (actin filaments, microtubules), actomyosin contractions, and endoplasmic reticulum calcium channels were essential for remote fiber inductions, while membrane mechanosensitive integrin and Piezo showed coordinative role in regulating the fiber assembly. The study provides new insights on cell mechanics-induced isotropic matrix modeling with dynamic large-spatial scales and the associated cellular mechanism. The assembled biomechanical scaffolds with pre-designs may lead to applications in micro-tissue engineering. This work implicates heterogeneous tissue structures maybe partially derived from isotropic cell mechanics.

## Introduction

Tissues and organs in the body often have isotropic structures or shapes, which developmental process is coordinated by cells and extracellular matrix (ECM) modeling(Alford et al., 2015; Kleinman et al., 2003). Abnormal ECM modeling is a sign for disorders, such as fibrosis in liver and lung, cartilage degradation in joints, and ECM-associated genetic diseases(Theocharis et al., 2019). Several studies demonstrated tumor cells able to reconstruct ECM environments in facilitating the metastasis(Erdogan et al., 2017; Han et al., 2016; Pankova et al., 2016). Cells also show the capability in remote ECM modeling, for example, two explanted fibroblast tissues could build fiber bundles across 1.5-4 cm distance on the collagen hydrogel(Harris et al., 1981; Stopak and Harris, 1982). Hence, cells can format specific ECM structure which in return mediates the physiological outcomes of cells and tissues. Besides the chemical cues, the biomechanical aspect how cells and ECM coordinate at the large-spatial scale during morphogenesis and for tissue engineering has been an interesting topic to explore.

Cell mechanical communications are recognized from in vitro and in vivo studies(Alisafaei et al., 2021; Long et al., 2022). Some different from traditional chemical cues, mechanical communications between cells show the features with long-range or large-spatial transmission, precise direction, and functional outcomes at tissue scale. For example, cells are able to sense the positions of neighboring cells through compliant substrates(Reinhart-King et al., 2008; Winer et al., 2009). Long-range force enables assembly of mammary tissue patterns and collagen fibrillary remodeling(Guo et al., 2012; Shi et al., 2014). Cardiomyocytes achieved similar beating frequency when seeding on elastic substrate(Nitsan et al., 2016; Sapir and Tzlil, 2017). During *Xenopus* embryonic development, the cell cluster of neuronal crest moves collectively to the destination with supracellular contraction at the rear to assist directional migration(Shellard and Mayor, 2021; Shellard et al., 2018). These increasing evidences have supported distant mechanical communication as a common nature of cells.

The cellular mechanism on mechanical communications has also been under study. As shown from recent work including ours, cells have the capability to sense the traction force or substrate deformation transmitted from neighboring cells or induced by external probe, which provides a bridge in understanding cell-cell distance interactions(Ouyang et al., 2020; Zarkoob et al., 2018). At the molecular level, the mechanosensitive receptor integrin and ion channel Piezo display importance in remote attractions between two types of cells(Liu et al., 2020; Pakshir et al., 2019). The calcium channels on endoplasmic reticulum (ER) are indispensable for cell distant mechanical interactions(Ouyang et al., 2022). There is also long-range force transmission through cell-cell junctions during the collective migration of epithelial sheet(Aoki et al., 2017; Sunyer et al., 2016). So far, some important mechanism and signals have been identified for the mechanical communications, whereas the general map at this aspect is still limited.

As one interesting observation, cells are able to reconstruct the COL gel to generate fibrous bundles(Barnes et al., 2014; Stopak and Harris, 1982), which biophysical mechanism has attracted research attentions. Physical force from tissues is able to rearrange COL patterns with long-range order(Stopak et al., 1985). A later study confirmed the local movements of COL driven by fibroblast contractions leading to the fibrillary organization(Sawhney and Howard, 2002). In recent decade, more mechanistic work has emerged on the cells-driven fibrous morphogenesis. Finite element (FE) simulations predicted the alignment of COL gel induced by long-range biomechanical transmission, although the results were conditional with anisotropic cell shape and within cell pair(Abhilash et al., 2014; Wang et al., 2014). FE simulations combined with fibrous continuum modeling further yielded the measurements of cell traction force propagation(Hall et al., 2016). Application of optical tweezers to pull beads on the COL hydrogel revealed the long-range stiffness gradients surrounding the cell(Han et al., 2018). With synthetic fibrous materials with controllable biomechanical architectures, cell contraction induces the alignment of fibers(Baker et al., 2015; Davidson et al., 2020). In mimicking in vivo condition, experimental stretch or fluidic shear stress is also able to align long-range order of extracellular matrix (ECM)(Cassereau et al., 2015; Wu et al., 2017). Recent evidence shows that cells-reorganized fiber bundles bear the major tensile force to guide cell distant interaction and correlated migration(Fan et al., 2021). In general, how to understand cells-induced ECM remodeling at large-spatial or tissue scale is still challenging.

Apparently, cells conduct the work in remodeling ECM microenvironment in physiology and pathology, which intracellular signaling is to be understood. Cancer-associated fibroblasts (CAFs) can induce COL cross-link switch, and align fibrous COL or fibronectin to promote fibroblast long-range order or direct cancer cell migration(Erdogan et al., 2017; Li et al., 2017; Pankova et al., 2016). Early-stage cancer cell spheroids embedded into COL containing fibroblasts generated force for fibers alignment and CAFs induction(Jung et al., 2020). Our work also showed that motile kidney epithelial cells (MDCK) can actively recruit soluble ECM from the medium to assemble base membrane or COL architectures in directing spherical or tubular tissue formations(Ouyang et al., 2021; Wang et al., 2020). The cell signals of integrin, RhoA, and actomyosin contraction show importance in cells-mediated ECM remodeling(Brownfield et al., 2013; Erdogan et al., 2017). However, the fundamental molecular mechanism in cells remotely driving the modeling is largely unknown.

In this work, we tried to study the COL fibrillary remodeling induced by cell long-range traction force, as well as the involved mechano-signaling mechanism in cells. The research designs were based on large-spatial arrays of cell clusters and incorporated an important feature with dynamic contraction in simulations. This helped overcome the limitations of previously conditional studies, such as single or a few cells/clusters, a demand for polarize cell shape, or static traction force(Aghvami et al., 2013). In previous work, we observed the traction force from live cells is a dynamic factor instead of static through the hydrogel, which was derived from the highly motile cell clusters under constantly rotating (Guo et al., 2012; Ouyang et al., 2021), as reported by Bissel group as well(Tanner et al., 2012). Here, with incorporated dynamic tractions, the simulations with Molecular Dynamics yielded highly matching outcomes with observed COL fiber bundles from the experiments, which didn’t have conditional spatial-scale limitation. The designed single polygons with variable geometries led to well predicted structures. The cell cytoskeletal integrity, actomyosin contractions, and ER calcium channels were essential for the remote induction of COL fibers, whereas the role of membrane mechanosensitive integrin β1 or Piezo1 alone was less critical.

## Materials and Methods

### Cell Culture and siRNA transfection

Rat airway smooth muscle (ASM) cells were isolated from the tracheal tissue of 6-8-week old female Sprague Dawley rats as described previously(Duan et al., 2016). The cells were cultured in low sugar DMEM (Invitrogen) supplemented with 10% fetal bovine serum (GIBCO) and penicillin/Streptomycin antibiotics (GIBCO). The cells used in the experiments were generally maintained within 10 passages.

To perform siRNA transfections, ASM cells were seeded into 6-well plate at ~40% density, and the next day, rat ITGB1 siRNA (Thermo), Piezo1 siRNA (Horizon Discovery), or control siRNA (50 nM in 2 mL medium) was transfected into the cells by using 5 µL Lipofectamine 3000™ reagent. After incubation for about 8 h, changed the medium and waited for 48~72 h before further experiments.

### Preparation of PDMS molds

The micro-patterned arrays of cell clusters were prepared according to the method derived from our previous work(Guo et al., 2012), but with more variably geometrical distributions. Basically, we used the software AutoCAD to design the arrays of circle distributions, which circles were set as 200 μm in both diameter and depth, and 600-800 μm in mutual spatial separation between the centers. The parameters for different geometrical patterns are listed in Table 1. The silicon wafers printed with the designs were manufactured by Suzhou MicroFlu Technology Co. and In Situ Co., Ltd.

**Table 1.**
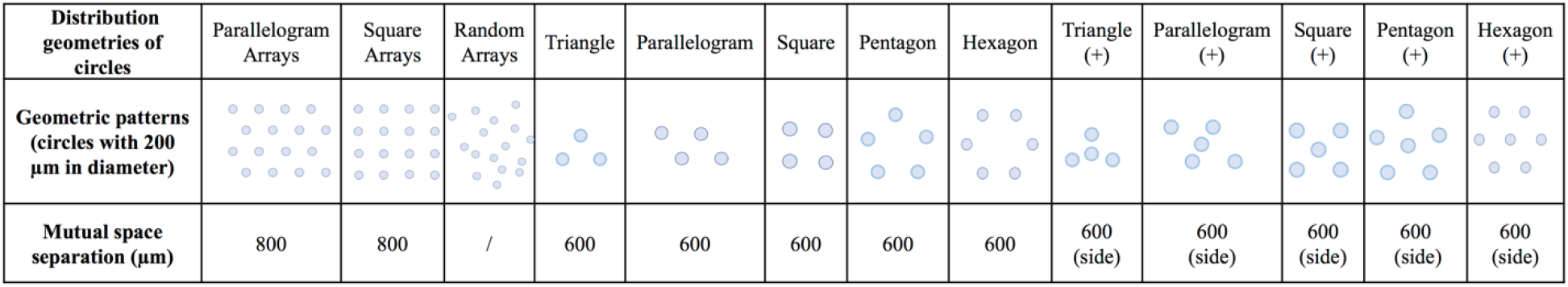
The parameters for variable geometric patterns.

| Distribution geometries of circles | Parallelogram Arrays | Square Arrays | Random Arrays | Triangle | Parallelogram | Square | Pentagon | Hexagon | Triangle (+) | Parallelogram (+) | Square (+) | Pentagon (+) | Hexagon (+) |
| --- | --- | --- | --- | --- | --- | --- | --- | --- | --- | --- | --- | --- | --- |
| Geometric patterns (circles with 200 μm in diameter) | | | | | | | | | | | | | |
| Mutual space separation (μm) | 800 | 800 | / | 600 | 600 | 600 | 600 | 600 | 600 (side) | 600 (side) | 600 (side) | 600 (side) | 600 (side) |

To fabricate the polydimethylsiloxane (PDMS) molds, the two liquid components from the Sylgard 184 Kit (Dow Corning) were mixed at a mass ratio of 10:1 (base to catalyst 87-RC), and poured onto the silicon wafer after extraction of air bubbles by vacuum, then cured at 80°C for over 4 h and left at room temperature overnight. The solidified PDMS was carefully peeled off from the wafer, and further cut into small pieces (~ 1×1 cm) to adapt the next experimental utility. The prepared PDMS molds were soaked in 75% ethanol for disinfection, and exposed to UV light for several hours in the cell culture hood before each experiment.

### Expression and purification of EGFP-CNA35 protein

The pET28a-EGFP-CNA35 DNA construct purchased from Addgene was described before(Aper et al., 2014). The procedures for expression and purification of EGFP-CNA35 protein were introduced in our recent work(Ouyang et al., 2020). In brief, the DNA plasmid was transformed into BL21 (DE3) competent bacteria, and when culturing in the Lysogeny Broth (LB) medium, the protein expression was induced by isopropyl β-D-1-thiogalactopyranoside (IPTG) at room temperature. The precipitated bacteria were lysed with B-PER lysate solution (Thermo Scientific), and HisPur Ni-NTA agarose beads were added to pull down the 6xHis-tagged protein. The concentration of purified protein was measured by Branford protein assay kit (Thermo), and protein aliquots were stored at -80°C.

### PDMS mold-printing onto agarose gel, and cell cluster culture

First, to prepare cell clusters on micro-patterned agarose gel. Agarose solution (20 g/L) was made by dissolving 2 g agarose (Thermo) in 100 mL sterile Phosphate-Buffered Saline (PBS) by microwaving, and ~150 μL agarose solution was dropped onto the glass-bottom dish. The sterilized PDMS mold was placed onto the solution, and slightly pressed downward. After solidification at room temperature, carefully peeled off the mold without disruption of the printed patterns on agarose gel. 200 µL suspension of ASM cells in medium (~1.2 × 10^6^ cells/mL) was added onto the agarose gel, and after 10-15 min waiting for cells to settle down into the pattern areas, absorbed the additional medium away and washed the patterned regions once or twice with culture medium carefully. Then, cells were generally filled into the circular caves (200 μm in depth) without much left on other areas (cells are non-adhesive on agarose gel).

Second, to cover the cell clusters with fluorescent COL (Cultrex 3-D Culture Matrix Rat Collagen I) hydrogel. The preparation of COL solution was carried out on ice: completely mixed the rat tail COL solution (4 mg/ml) with purified EGFP-CNA35 protein (diluted to 1 mg/ml in advance) at the volume ratio of 5:1, and waited for about 15 minutes; afterwards, added neutralizing solution at 1:9 volume ratio of the COL one, and further diluted the COL to final 1 mg/ml (or other alternative concentration) with PBS. Then added the mixed COL solution (300-400 μL) onto the patterned cells-containing agarose gel, and tried to spread the COL liquid across the glass-bottom surface of the dish in addressing none adhesive COL gel onto agarose. Then placed the assembled cell samples in the culture incubator at 37°C for 15~20 min, and after gel, added 2 ml of regular culture medium into the dish. In this way, the cell clusters were generally embedded into COL while underneath the gel layer.

### Cell cluster cultures along with inhibitor applications

Before detached for generating cell cluster arrays, ASM cells cultured in six-well plates (NEST) were pre-incubated with the inhibitors for 1~2 h, and the solvent DMSO (Beyotime) was used as control. After cell clusters were sandwiched between COL and agarose gels, the culture medium containing the same inhibitor was added back. To enhance the medium diffusion with inhibitor through the hydrogel, a small hole was created at the edge of the gel with a fine needle without disturbing the cell mass. After culture for 24 hours, 1 mL more medium with the same inhibitor was further supplemented to the culture in considering the inhibitor stability at 37°C. The formation of COL fiber bands and cell migration were recorded under microscopy.

The applied inhibitors in the experiments were mostly purchased from Sigma, including 2-Amino-ethoxydiphenyl borate (2-APB, an IP_3_R calcium channel inhibitor with cell permeability, 20 µM), Nifedipine (membrane L-type calcium channel inhibitor, 20 µM), Lanthanoum(III) chloride (LaCl_3_, SOC calcium channel inhibitor, 100 µM), Nocodazole (microtubule de - polymerizing agent, 1 µM), Blebbistatin (myosin II ATPase inhibitor, 40 µM), Cytochalasin D (CytoD, actin polymerization inhibitor, 1 µM), ML-7 (MLCK inhibitor, 40 µM), NSC23766 (Rac1 activation inhibitor, 100 µM), and Latrunculin A (LatA, microfilament polymerization inhibitor, 237 nM). Y27632 (ROCK inhibitor, 40 µM) was from MedChemExpress, and Thapsigargin (ER SERCA pump inhibitor, 10 µM) from Abcam.

### Q-PCR measurements

After ITGB1 or Piezo1 siRNA transfection, the decrease of mRNA expression in ASM cells was evaluated by real-time quantitative PCR (qPCR), which details were described in our recent work(Fang et al., 2022). The primer sequences of qPCR derived from others’ studies are listed on Table 2(Beca et al., 2021; Wang et al., 2019).

**Table 2.**
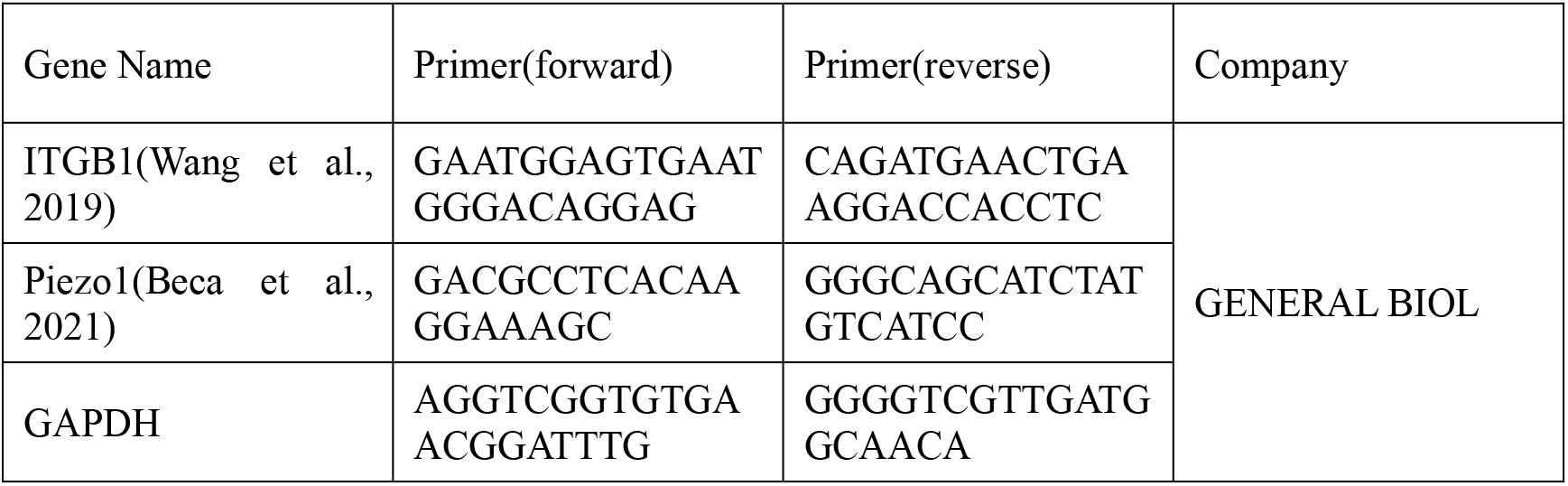
The primer sequences of qPCR for measuring ITGB1 and Piezo1 mRNA levels.

### Molecular dynamics (MD) simulations of fibers clustering and reconstruction

A 2D network model based on MD simulation was introduced to investigate the mechanics of fibers cluster and cell migrations. The collagen fibers were simplified to bonds on a plane, and their crosslinks were set as the nodes between different bonds. The collagen fiber is an isotropic elastic material that has a high modulus in tension and a very low modulus in compression(Ushiki, 2002). The bond’s tension stiffness was set as 100 times of compression stiffness. The fibers crosslinks may reconstruct under tension or compression. Here we set a reconstruction judgmental condition that bond length change is higher than 50 percent of bond length. The crosslink of this fiber will be reconstructed and this bond length will change from its initial value to its current value. The fibers network under cells region were removed, and cells were shown as a circle with 83 nodes. The Stochastic dynamics (SD) were performed by GROMACS package(Wennberg et al., 2015). The XYZ coordinates of cells nodes and Z coordinates of crosslinks nodes were fixed during simulation. The dynamic contraction of cells was applied as the radius reduction of the cells’ circles. The circle radius was reduced by 5% per 50000 simulation steps. The simulation results were analyzed with VMD program(Humphrey et al., 1996).

### Finite element modeling (FEM) for stress distributions on the hydrogel

The FEM model to investigate the correlation between matrix stress distribution and the fibers network formation has been introduced in our previous studies(He et al., 2015; Ouyang et al., 2020). The interaction surface of cells and collagen matrix was simplified to an isotropic linear elastic plane surface. The cell region was removed, and cell traction force was applied on the edges of cells’ region to mimic the contraction force of cells on the matrix surface. The stress value on the matrix was normalized to reach a relative quantitative analysis and results. The FEM model was solved and visualized by ABAQUS packages.

### Live cell imaging, and quantifications of COL fiber intensities

The live cell microscopy workstation (Zeiss) was equipped with an X-Y-Z controller system for multi-position imaging, fine auto-focusing function, and a culture chamber loaded on the sample stage to maintain 5% CO_2_ and the temperature at 37°C. Most fluorescence imaging experiments were carried out with x5 and x10 objectives under the microscope. One round of the experiments spanned 2-3 days, and the images for fluorescent collagen fibers and cell migrations were taken at different time points as indicated, for example, the initial status, 12, 24, 36, 48 h.

ImageJ was used to quantify the relative fluorescence intensities (FL) of collagen fiber bands in contrast to the close non-fiber areas, as demonstrated in Figure S1. In the procedures, the image was rotated so the COL fibers were displayed in horizontal direction; a rectangular region (ROI) with same size was chosen in covering each COL fiber band and its close area while the fiber band was positioned at the middle of ROI. Averaged fluorescence intensity (FL) was measured from left toward right along the selected rectangular ROI by using the “plot profile” function in ImageJ. The acquired values from each ROI were input into the Excel file, which generated an averaged fluorescence distribution curve along the selected region. Then, the sum (S_max_) of seven points around the maximum intensity value (three values from the left and another three from the right) was calculated, as well as the sum (S_min_) of seven values from two ends of the curve (three or four values from each) in representing the low non-fiber intensities. The FL ratio was defined as S_max_ divided by S_min_ (Ratio = S_max_/S_min_). Generally, fluorescent COL images were quantified from 4 or 5 time points within 48 h, for instance, 0, 12, 24, 36, 48 h. Statistical comparison was conducted between the initial time point and later ones, which presents the emerging COL fibers induced by cell clusters remotely.

The COL fluorescence curves and graphs of scattering dots were processed by Origin 2020, GraphPad Prism 6, and Excel software. Student’s t-test was applied for statistical analysis. *, **, ***, **** represent ‘p value’ < 0.05, 0.01, 0.001, 0.0001 for significant difference respectively, while ‘NS’ for no significant difference.

## Results

### Molecular Dynamics and Finite Element simulations for COL fiber clustering driven by cell stress distribution

To study how cell traction force remotely drives clustering of COL fibers on the hydrogel, we built a 2D network model based on MD simulation to investigate the direction and mechanics of fibers cluster (Fig.1A). The collagen fibers were simplified to bonds on a plane, and their crosslinks were set as the nodes between different bonds (Fig.1B). The collagen fiber is an isotropic elastic material that has a high modulus in tension and a very low modulus in compression (Fig.1C) (Ushiki, 2002).

**Figure 1.**
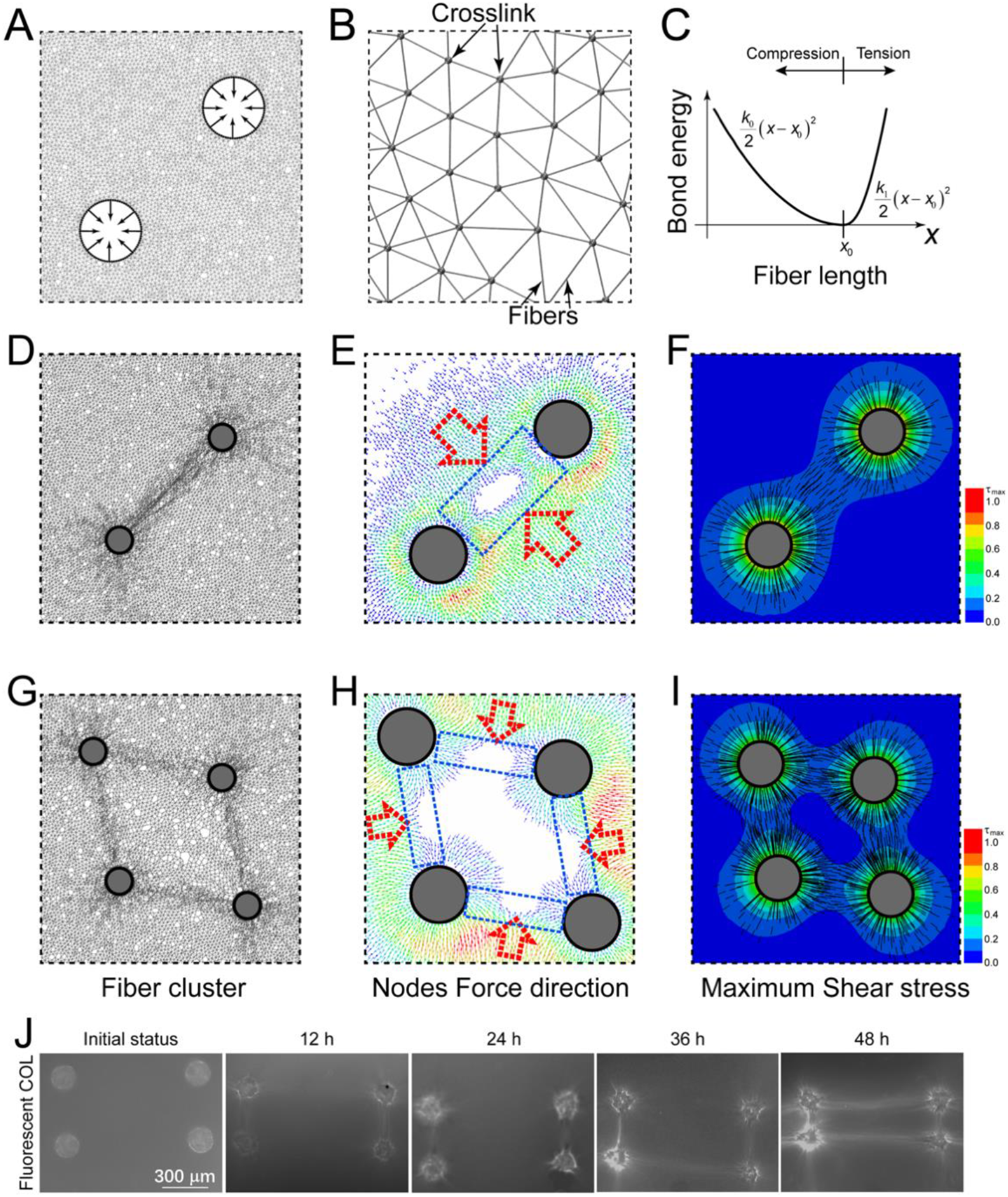
The corresponding relation between fibers clustering and stress distribution. **(A)** The molecular dynamics simulation model for fibers clustering. The collagen fibers were simplified to a bonds network. The contraction of cells was applied on the boundary of the empty regions. **(B)** A detail view of the fibers bonds network. **(C)** The bond energy changes with bond length. When bond was compressed or tensed, the tension spring stiffness is higher than compression spring stiffness. **(D)** Fibers were recruited in the connection line of two cells. **(E)** The particle force distribution and the directions of particle force in two cells by MD simulation. **(F)** The maximum shearing stress distribution on matrix and the directions of first principle stress in two cells condition. **(G)** Fibers were recruited in the connection line of relative close cell pairs in the condition of four cells. **(H)** The particle force distribution and the directions of particle force in four cells by MD simulation. The colors from blue to red represent lower to higher simulated stress in (E) and (H). **(I)** The maximum shearing stress distribution on matrix and the directions of first principle stress in four cells condition. **(J)** The representative experimental result of fluorescent COL fiber clustering induced by cell clusters at the indicated culture time. The center distances of the paired clusters and between two pairs are 500 and 1000 μm.

To investigate how stress distribution regulates fibers clustering, the force direction on each node and maximum shearing stress distribution on matrix was calculated by molecular simulation and FEM respectively. As shown in Fig. 1D, two cells shrunk with dynamic contractions, and fibers cluster between cells from MD simulation. The particle force in MD also revealed similar results (Fig. 1E). Considering the orientation of particle force, the fiber was bearing a pulling force toward the connection line between cells, which would lead to the fibers clustering (seen in Fig. 1D&E). The local maximum shearing stress distribution on matrix was calculated and shown in Fig. 1F, and the black line revealed the direction of first principle stress at local place. The maximum shearing stress has a relatively high value around the connection line of two cells, and this region is consistent with the fibers clustering (Fig. 1F).

For the condition of four cells, fibers clustering formed in relatively closer cell pairs instead of central region of four cells (seen in Fig. 1G). Similar as the condition of two cells, the fiber lines are also in agreement with the first principle stress direction and the particle force distributions (Fig. 1H). The central region has a higher first and second principle stress, but the maximum shearing stress is lower than the connecting lines of cell pairs (Fig. 1I). Experiments with seeded cell cluster pairs demonstrated the COL fibers emerging between clusters (Fig. 1J), consistent with the MD simulated outcomes. Taken together, fibers were recruited in the region with higher maximum shearing stress and the orientation of local Maximum Principle Stress.

### Large-scale COL fiber remodeling induced by geometrical square and parallelogram-arranged cell cluster arrays

The simulated outcomes by Molecular Dynamics (MD) showed fiber cluster occurrence mostly at the sides based on the dynamic contraction in geometrical square-positioned cell cluster arrays (Fig. 2A). The experimental results were acquired by seeding cell clusters into square or parallelogram arrays covered with fluorescent COL gel (1 mg/ml, labeled with EGFP-CNA35), and images were taken at indicated time points in the next two days. The data showed the emerging square lattices of COL fiber bundles, resulting into connection of the cluster array (Fig. 2B). More details with the gradual fiber growth were shown in Supplementary Information (Fig. S2A). The fluorescence quantification (Ratio = S_max_/S_min_) and intensity distributions indicated the gradual growth of the fibers (Fig. 2C, D). The ratio started at ~1.0 as an initial uniform fluorescence distribution, and reached ~1.2-1.5 as fiber growth between cell clusters. Fluorescence curves of individual selected ROI regions are presented in Fig. S3. The fiber lattice structure stayed stable in the two-day observations, indicating balanced contraction force through the square array.

**Figure 2.**
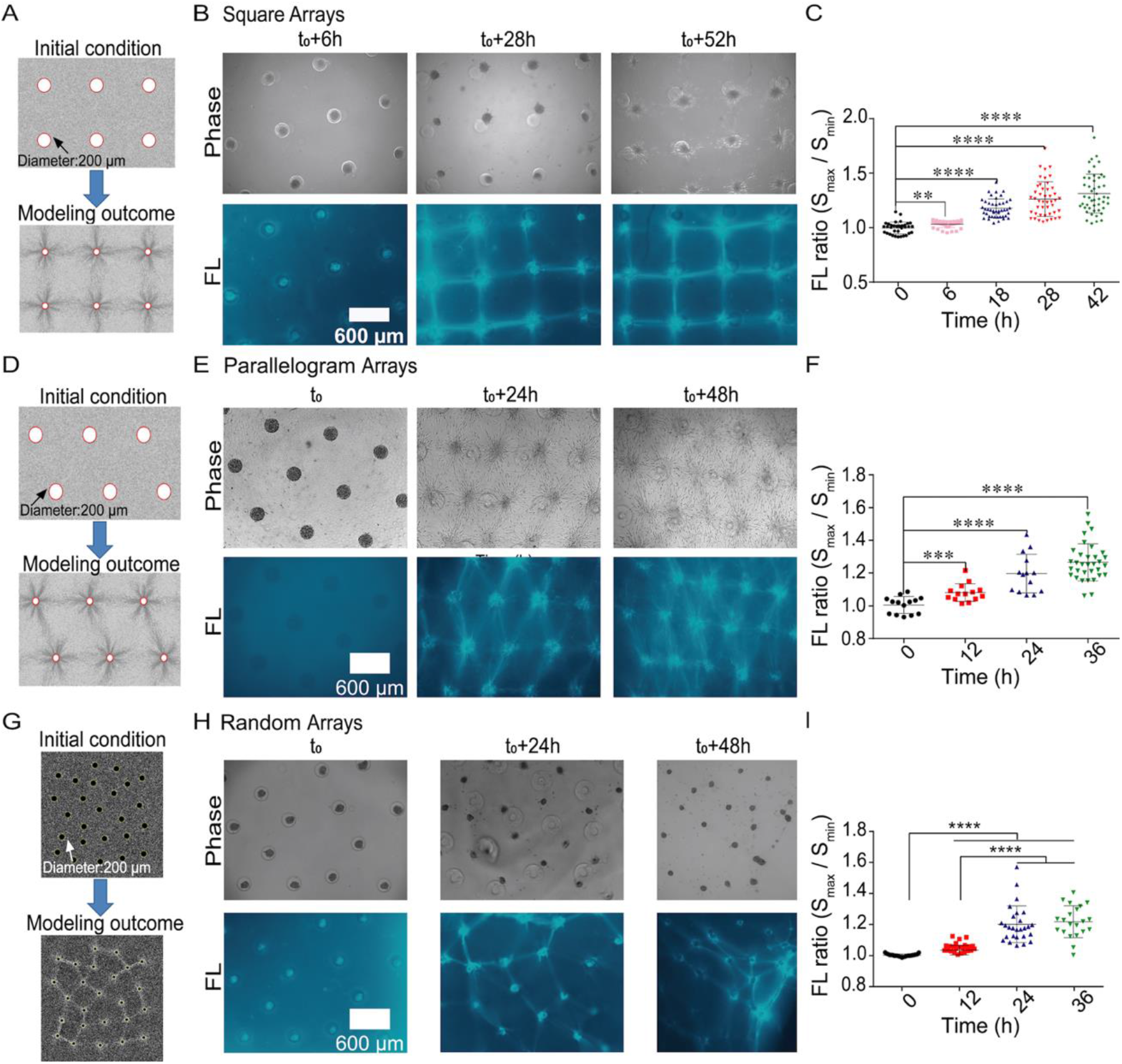
Remote COL fiber inductions by variable arrays of cell clusters. ASM cell clusters were seeded into the designed geometrical arrays with fluorescent COL hydrogel (1 mg/ml) atop. Images of cells and COL fluorescence were taken at the indicated time. **(A)** MD simulated outcome of square arrays based on dynamic contraction force of cell clusters. **(B)** The emerging COL fiber bundles between cell clusters positioned into square arrays. **(C)** Fluorescence (FL) quantification of COL fibers (mean ± S.E.M.) in contrast to the non-fiber region based on the ratio calibrations (S_max_/S_min_). Each dot represents the ratio value of one fiber bundle. **(D)** The spatial distributions of averaged COL fluorescence along with the selected regions (300 μm in width). The quantification process is described in the Methods. **(E-H)** The MD simulation outcome (E), the emerging COL fiber bundles (F), FL ratio quantifications (G), and fluorescence distributions (H) from parallelogram-arranged cell cluster arrays. **(I-K)** The MD simulation outcome (I), the emerging COL fiber bundles (J), FL ratio quantification (K) from the geometrical random-style cluster arrays. The statistical significance for COL fluorescence between the initial and indicated time points was evaluated by Student’s t-test. *, **, ***, **** represent p value < 0.05, 0.01, 0.001, 0.0001, respectively, and so on through the paper.

We next examined the geometrical parallelogram arrays. MD simulation showed fiber clustering at the sides while less obvious at the diagonal directions (Fig. 2E). When cell clusters were seeded into parallelogram array mode, COL fiber buckling occurred mostly at the sides, resulting into formation of parallelogram lattices (Fig. 2F, more details shown in Fig. S2B). Fluorescence quantification and spatial distributions indicated the gradual accumulations of COL fibers between cell clusters (Figs. 2G&H, S3B). As seen at later time (e.g. 36 h) (Fig. S2A, B), certain fiber bundles showed up at the diagonal directions in parallelogram and square arrays. This experimental difference from the simulations might be partially due to the changed material property under remodeling by the cell clusters, which explanation demands further study.

We further applied random-style distributed arrays of cell clusters (mutual spatial separations within 600-1000 μm) to check the simulation and experimental outcomes for fiber growth. The MD simulation well turned out fiber emergences in connecting cell clusters (Fig. 2I), which dynamic process is shown in Movie 1. Experiments based on random-positioned arrays showed gradual fiber growth, resulting into fiber network formation along with clusters (less stable structure) (Figs. 2J, K and S2C), which indicates the robustness of cell pattern-induced fiber assembly. Therefore, the experimental results of fiber clustering within variable geometries stayed pretty consistent with MD and FEM simulated outcomes based on dynamic tractions at the large-spatial scale.

### COL fiber assembly induced in single polygons with diverse geometries

We further designed cell clusters positioned into single polygons including triangles, parallelograms, squares, pentagons, and hexagons (600 μm in mutual separations of the centers). This provided the chance to assess the robustness of cell contraction-induced fiber assembly, and the stability of assembled fiber structures.

When cell clusters were settled down into single triangles, strong COL fibers were shown up at the three sides rapidly (within 12 h), and cells migrated toward each other (Fig. 3A, left). There was also fiber growth between two triangles which had 1000 μm in distance. Group fluorescence quantification confirmed fiber assembly in the designed triangles (Fig. 3A, right). In single parallelograms, COL fibers emerged gradually at the four sides along with some shown up at the diagonal lines (600 μm in distance) (Fig. 3B). In single squares, COL fibers occurred at the sides, not much at the diagonal direction (848 μm in distance) (Fig. 3C). When cell clusters were arranged into single pentagons or hexagons, COL fibers were shown up efficiently at the sides, but not at the diagonal lines (Fig. 3D, E), which distances between cell clusters at the diagonal directions (~1141, 1039 μm in pentagons, or 1200 μm in hexagons) were much larger that at the sides (600 μm). Cells also migrated toward each other along the sides.

**Figure 3.**
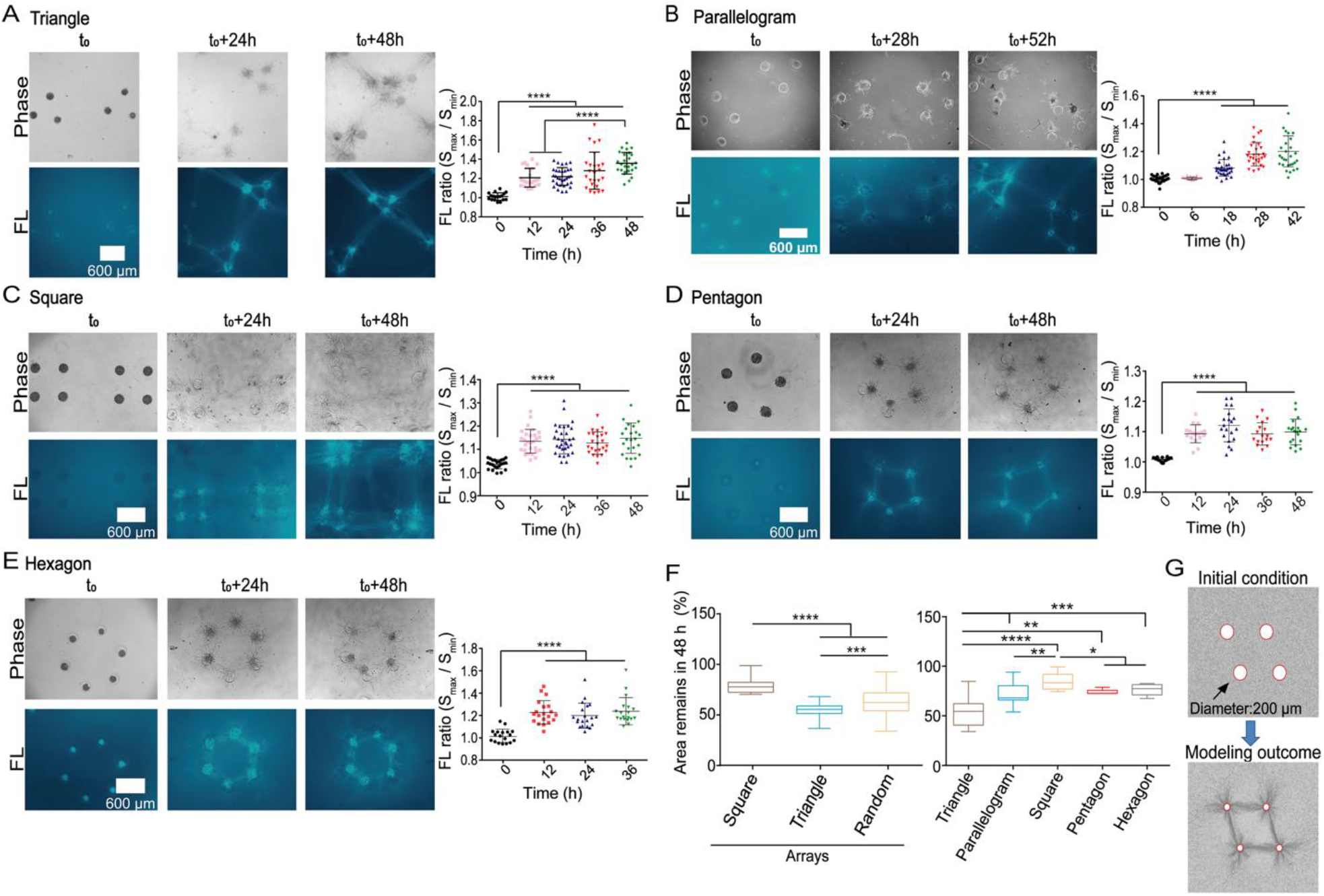
Fiber assembly in single polygons with diverse geometries, and their structural stabilities. Cell clusters were seeded into single polygons. Images were acquired at different time points, and fluorescent COL fibers were measured as Ratio = S_max_/S_min_. **(A-E)** The growth of COL fibers, and fluorescence quantification (mean ± S.E.M.) for fiber strength in single triangles (A), parallelograms (B), squares (C), pentagons (D), and hexagons (E). **(F)** Stability measurements with remained areas (%) from geometrical shrinking for the diverse arrays (from Figure 2) and single polygons within 48 h. The mean ± S.E.M. values for square array: 79.5±1.3; parallelogram array: 55.3±0.7; random array: 63.3±1.8; single triangle: 54.8±3.5; parallelogram: 71.6±2.6; square: 82.4±2.4; pentagon: 74.3±1.2; hexagon: 75.2±1.9. (**G**) The MD simulation for fiber clustering in a single parallelogram.

By measuring the area changes within 48 h culture to evaluate the structural stabilities shown in Figure 2, the square arrays were more stable than parallelogram and random-style ones (Fig. 3F, left). The single polygons (Fig. 3A-E) were quite stable (less than 30% shrinking in areas), except triangle ones (~45% shrinking) (Fig. 3F, right). As a demonstration, MD simulation for single parallelogram predicted fiber clustering mostly at the sides (Fig. 3G), and similar to FEM simulation of the maximum stress distribution (Fig. S4A). The more details for fiber inductions from single triangles to hexagons are presented in Figure S4(B-F). Together, these results from above confirmed the high consistency between the dynamic simulations and experimental outcomes (especially at early time stages), and demonstrated fiber assembly induced by cell remote mechanics in COL hydrogel.

### The cytoskeleton integrity and actomyosin contraction are essential for remote induction of COL fiber assembly

Since cells induced fiber assembly in the experimental models, we further looked at the involved cellular biomechanical mechanism by applications of the large-spatial parallelogram arrays. First by checking the importance of cell cytoskeletons, cells actively migrated out from clusters towards each other, and well developed COL fiber networks at control condition (0.1% DMSO) (Fig. 4A); when actin cytoskeleton was inhibited with cytochalasin D (CytoD) or latruculin A (LatA) treatment, the fiber assembly was gone, and cells didn’t migrate out actively (Figs. 4B&C, S5C); similar results were observed when microtubule cytoskeleton was inhibited with Nocodazole (Fig. 4D). The comparison of percentage changes in COL fluorescence quantification showed inhibited fiber assembly by the three drug treatments (Fig. 4E). The representative images with more details for cytoskeleton inhibitions are presented in Figure S5(A-F). In this work, cell migration was just descriptive but not quantitative, as not the main focus of the study. These data indicate that the integrity of both cellular actin and tubulin cytoskeletons are essential for remote induction of fiber assembly.

**Figure 4.**
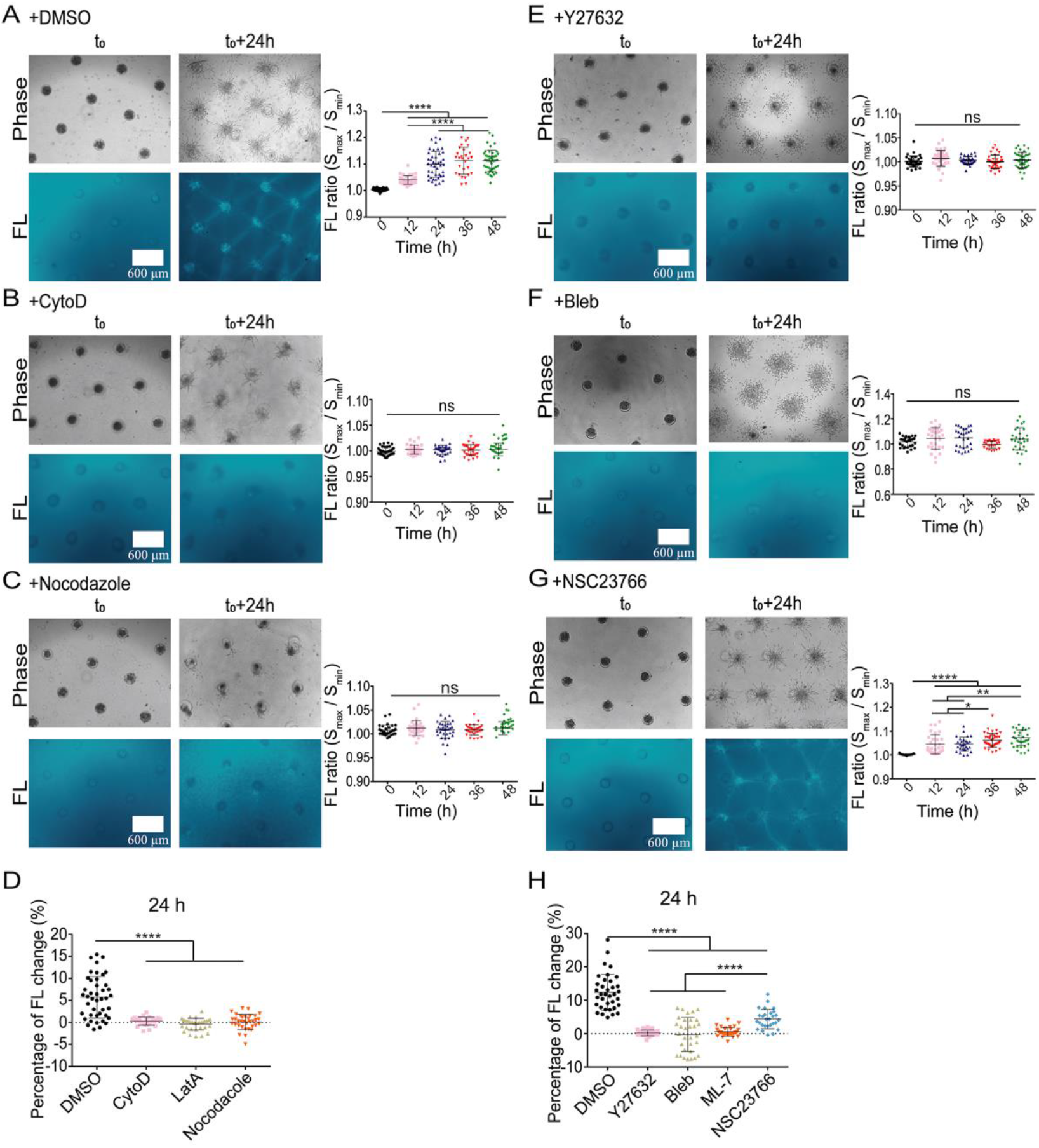
Cytoskeleton integrity and actomyosin contraction in remote induction of fiber assembly. Cell clusters were cultured into large parallelogram arrays with cytoskeleton or actomyosin inhibitors. **(A-C)** The COL fiber growth and fluorescence quantification at control condition (DMSO) (A), or under treatment with 1 µM CytoD (B), or 1 µM Nocodazole (C). **(D)** Comparison for the percentage changes of COL fiber fluorescence within 24 h with or without cytoskeleton inhibitors. **(E-G)** The COL fiber growth and fluorescence quantifications under treatment with 40 µM Y27632 (E), 40 µM Blebbistanin (F), or 40 µM NSC23766 (G). (**J**) Comparison for fiber fluorescence change (percentage) within 24 h at the indicated conditions. The experimental data with LatA or ML-7 treatment were shown in Figure S6.

We continued to check the role of cellular actomyosin contraction which is mainly regulated by RhoA-ROCK-Myosin II signaling pathway(Vicente-Manzanares et al., 2009). In contrast to the control group, the induced fiber assembly disappeared after inhibition of ROCK with Y27632 (Fig. 4G). When inhibited actomyosin contraction with myosin ATPase inhibitor Blebbistanin, or myosin light-chain kinase inhibitor ML-7, fiber assembly was also blocked (Fig. 4H, I). The images with more details are presented in Figure S6(A-E, H). After the contraction inhibition, cell migrations were significantly reduced and shown in a scattering way but not directional (Fig. 4G-I). When inhibition of small GTPase Rac1 with NSC23766 (40 μM), fiber assembly showed up with partial reductions, and cell migrations were less active (Figs. 4J, S5F). Similar results were observed with more reduced cell migrations at higher concentration of NSC23766 (100 μM) (Fig. S6G). The comparisons from fluorescent quantifications showed blocked fiber assembly after the contraction inhibitions, while partial inhibition with Rac1 inhibitor (Fig. 4K). These data together indicate that actomyosin contractions from cell clusters are essential for remote induction of COL fiber assembly.

### Calcium channels mediate cell clusters-induced fiber assembly

Our recent work showed that calcium channels at endoplasmic reticulum (ER) membrane are critical in cell-cell mechanical communications(Ouyang et al., 2022), hence the importance in the remote induction of fiber assembly was examined here. Similarly, COL fibers were assembled into network structure along with active cell migrations under control condition in the parallelogram arrays (Fig. 5A). When inhibition of inositol 1,4,5-trisphosphate receptor (IP_3_R) calcium channel or SERCA pump at ER membrane with 2-APB or Thapsigargin, both fiber assembly and cell migrations were almost blocked (Fig. 5B, C). Inhibition of L-type calcium channel at plasma membrane with Nifedipine also resulted into the large inhibitions (Fig. 5D), whereas inhibition of membrane Store-operated calcium channels (SOCs) with LaCl_3_ had little effect (Fig. 5E). The images with more details are shown in Figure S7(A-E). The percentage increases in fiber fluorescence quantification confirmed these observations (Fig. 5F). These results indicate that calcium signals regulated by the ER store and plasma membrane are important for cell clusters-induced fiber assembly.

**Figure 5.**
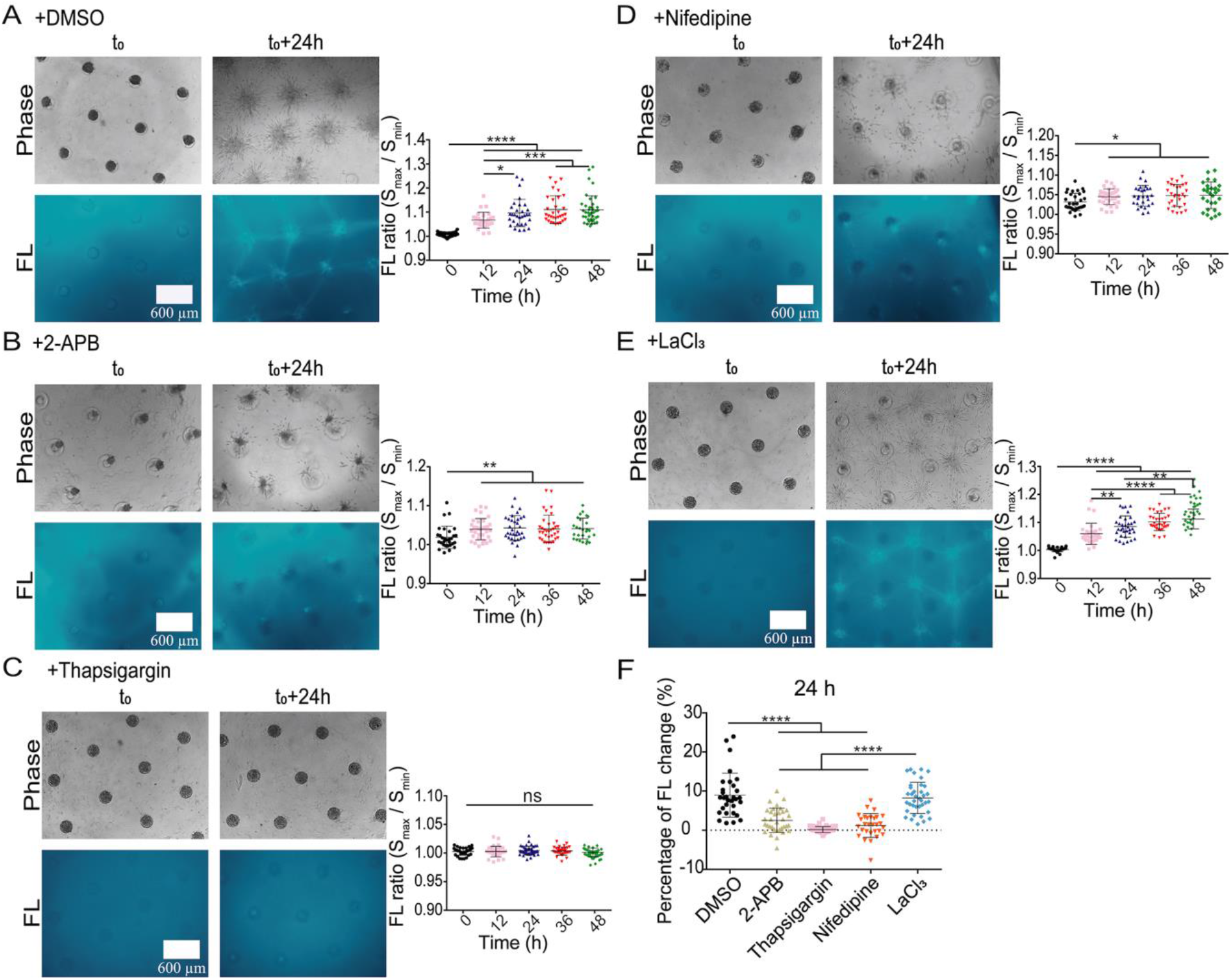
Calcium channels in regulating cell clusters-induced fiber assembly. Cell clusters were positioned into geometrical parallelogram arrays, and treated with calcium channel inhibitors. **(A-E)** Fiber assembly and fluorescence quantifications at control condition (DMSO) (A), or treated with 20 µM 2-APB (IP_3_R channel inhibitor) (B), 10 µM Thapasigargin (SERCA pump inhibitor) (C), 20 µM Nifedipine (L-type channel inhibitor) (D), or 100 µM LaCl_3_ (SOCs inhibitor) (E). **(F)** Comparison for fiber fluorescence changes (%) in 24 h under the indicated conditions.

### The importance of membrane mechanosensitive integrin β1 and Piezo in remote induction of fiber assembly

Plasma membrane integrin and Piezo1 are mechanosensitive components(Ouyang et al., 2022; Pakshir et al., 2019). Here, we examined their roles in the remote fiber assembly. After partial down-regulation of integrin β1 expression with siRNA transfection in ASM cells, COL fibers were partially induced and cell migration was less active in comparison to the control groups (Fig. S8A-C). Fluorescence quantification showed certain inhibition of fiber assembly with integrin β1 down-regulation alone (Fig. S8D&E). Then we further examined double down-regulations of integrin β1 and Piezo1 with siRNA transfections in which COL fiber inductions were significantly inhibited in comparison to the control group (Figs. 6A-C and S9A&B). These data indicate a possible coordinative role by integrin β1 and Piezo1 in the remote fiber induction, which is also supported by cell mechanosensing mechanism in recent work(Yao et al., 2022).

**Figure 6.**
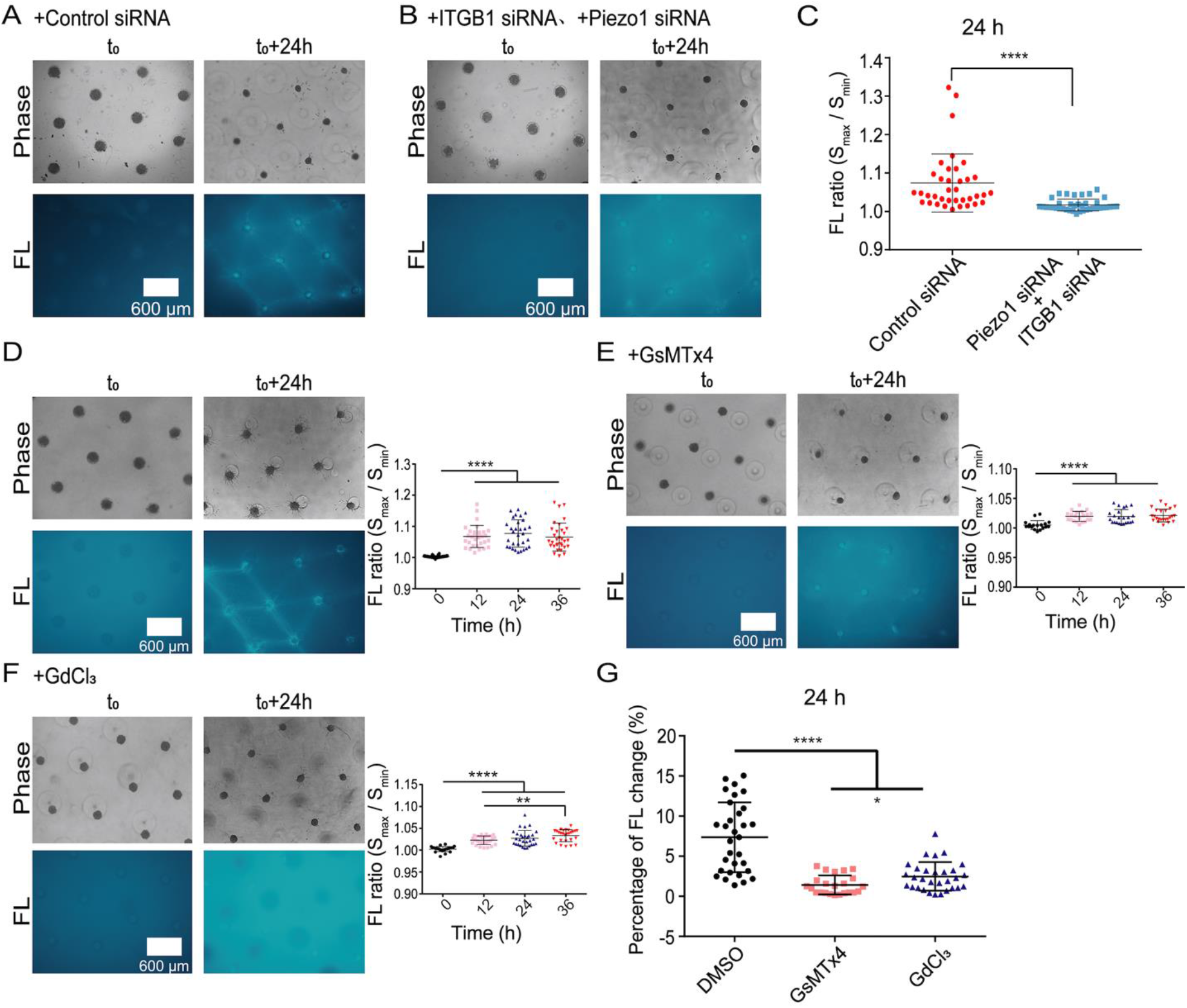
The role of integrin β1 and Piezo in fiber inductions. ASM cells were transfected with negative siRNA (100 nM) or double positive siRNA for integrin β1 (ITGB1, 50 nM) and Piezo1 (50 nM). COL fiber inductions were recorded and quantified in geometrical parallelogram-arrayed clusters. **(A-C)** Fluorescent COL fibers and quantifications under the condition of cell clusters transfected with negative siRNA or double ITGB1 and Piezo1 siRNA. **(D-F)** Fluorescent COL fiber induction and quantification under the control condition (D), or with treatment of 5 μM Piezo inhibitor GsMTx4 (E) or 100 μM GdCl_3_ (F). **(I, J)** The mRNA expression levels (I) and percentage changes in fiber growth (J) in control groups or Piezo1 siRNA-transfected group.

In considering that lung tissue expresses both Piezo1 and Piezo2(Lee et al., 2014), we further tested the chemical inhibitors of Piezos during the fiber induction by ASM cells. Two Piezo inhibitors GsMTx4 and GdCl_3_ both significantly reduced the COL fiber assembly in the large-spatial parallelogram cluster arrays (Figs. 6D-G). More details in images are shown in Fig. S10A-C. The data indicate that the two types of membrane mechanosensitive molecules integrin and Piezo together were critical in remote induction of fiber assembly by ASM cell clusters.

## Discussion

The long-range or large-spatial mechanical communications and resulted ECM modeling provide new angle in understanding physiological homeostasis in vivo, tissue morphogenesis, and the progressing in certain diseases. In this work, we tried to investigate the biophysical mechanism of large-spatial fiber assembly induced remotely by cell mechanics, and the supporting cell molecular mechanism. By incorporating dynamic contractions, Molecular Dynamics (MD) along with Finite Element (FE) modeling was able to simulate large-scale isotropic COL fiber clustering under variable geometrical distributions of cells. Experimental outcomes from cell clusters in spatial arrays yielded similar results to the simulated ones without spatial limitation, which also provided a platform in examining involved mechanical signals.

### Large-spatial and dynamic simulations for isotropic COL fiber clustering by MD and FE modeling along with cell experimental verifications

According to our previous observations that cells-derived traction force was dynamic but not static in the hydrogels, MD simulation containing the dynamic factor was applied to model the COL fiber clustering induced remotely by cells (Fig. 1A-C). The simulated results were consistent with the calibrated maximum shear stress distributions from FE modeling (Fig. 1D-I), which didn’t have pre-conditional limitations like cell shape, cell number, geometry, or space. To check whether the simulated results occurred in reality, we designed experiments with variable geometrical arrays for cell cluster distributions (Figs. 1J and 2). The geometrically arranged cell clusters including large-spatial square, parallelogram, and random-style arrays were covered into a layer of fluorescent COL hydrogel. The three types of arrays yielded highly consistent results with the MD simulated ones under similar setups (Fig. 2). These data largely verified the MD simulations based on cell dynamic shear tractions, and also demonstrated the robustness of ECM modeling from pre-designs.

### COL fiber inductions in single polygons, and their contraction-associated structural stability

Beside the large-scale arrays, it is also interesting to examine fiber inductions in single polygons with different geometries. The appropriately seeded cell clusters produced triangles, squares, parallelograms, pentagons, and hexagons through COL fiber remodeling (Fig. 3A-E). Due to the assembled fiber scaffolds induced by cell contractions, their structural stability is measured in area changes during 48 h culture. The large square array (~20% shrinkage) is more stable than parallelogram and random-style ones (~40-45% shrinking) (Fig. 3F, left graph). In single polygons, the squares are the most stable (~18% shrinking), followed with pentagons and hexagons (~25% shrinking), then parallelograms (~30% shrinking) and triangles (~45% shrinking) (Fig. 3F, right graph). The stability of the assembled scaffolds can reflect the internal contraction strength and balance.

This helps explain cell contractions-induced COL remodeling, and implicates the existence of tensions in maintaining stability of the fiber structures. It is consistent with recent report that the fibers bear the major tensions between cells(Fan et al., 2021). The MD simulations predicted the fiber clustering in single parallelogram (Fig. 3G), but not yet tension-mediated structural stabilities.

### The cellular mechanical mechanism in mediating the remote inductions of fiber assembly

Inhibitions of cytoskeletons (actin microfilaments and microtubules) or myosin contractility blocked the fiber inductions (Fig. 4), which demonstrates cell contractions-dependent remodeling of COL hydrogel. It is noted that tubulin cytoskeleton, which doesn’t provide cellular contraction force but gives mechanical supports, is also essential for the remote fiber inductions (Fig. 4D, E). Although there is no direct data to interpret this result yet, it may be related to microtubule functions in motility and intracellular transportations(Janke and Magiera, 2020).

Calcium signal is mechanosensitive in cells(Kim et al., 2015; Kim et al., 2009). In this work, inhibitions of ER calcium channels IP_3_R and SERCA pump, or plasma membrane L-type calcium channel largely reduced the capability of cell clusters in remote induction of fiber assembly (Fig. 5). Therefore, the physiological flow of Ca^2+^ between cells and the microenvironment, or between the cytoplasm and ER store is essential in driving the hydrogel remodeling. Besides other rich functions of calcium signals, in regarding the role of Ca^2+^ in regulating actomyosin contractility(Allen and Leinwand, 2002; Batters et al., 2016), it can make certain connection for these mechanosensitive calcium channels in the fiber inductions.

Studies have shown the importance for the membrane mechanosensitive integrin (β1) or Piezo1 in cell distant mechanical interactions(Liu et al., 2020; Ouyang et al., 2022; Pakshir et al., 2019). Here, we further examined the role in cell remote induction of fiber assembly. By applications of siRNA-transfection method or chemical inhibitors, integrin β1 and Piezo1showed coordinative roles in mediating the fiber assembly (Fig. 6A-G). Since the remote fiber assembly is induced from contraction of cell clusters, integrin may have the role in mechanical transmission between cells and the surrounding hydrogel.

Although cell migrations were not a major focus in this work, the cytoskeletons of microfilaments and microtubules seemed essential for cell movements in the experimental model (Fig. 4A-D), and myosin contraction and ER calcium channels are critical in cell directional migration between clusters (Fig. 4F-J).

### The prospective application for cultured scaffolds in tissue engineering

This work also yields one principal experience: to culture cells-induced scaffolds or tissue patterns according to pre-designs. Tissues and organs have specific shapes or structures, which has been one intriguing topic in developmental biology, but also a challenge in tissue engineering. Here, based on inductions of COL hydrogel remodeling with pre-designed cluster distributions, we successfully produced square/parallelogram/random-array lattices with assembled fibers (Fig. 2), and also single polygonal structures from triangles to hexagons (Fig. 3), although their structural stabilities were variable. This may lead to a new pathway for micro-tissue bioengineering including neural network, skin tissue, microvascular system with supplemental recipe in the current field.

## Conclusions

This work provided new mechanistic insights with dynamic and spatial factors on the isotropic induction of ECM modeling by cell remote mechanics, as well as the associated cellular mechanical mechanism. As depicted in Figure 7: by incorporated dynamic contractions, the simulations with Molecular Dynamics yielded highly matching outcomes without spatial limitation to isotropic COL fiber assembly in experiments from large square, parallelogram and random-style arrays of cell cluster cultures. The designed single polygons with variable geometries led to well predicted structures with assembled COL fibers. The cell cytoskeletal integrity (actin filaments and microtubules), actomyosin contractions, and ER calcium channels were essential for remote induction of COL fibers, while membrane mechanosensitive integrin β1 and Piezo1 show coordinative role in mediating the fiber assembly. The assembled scaffolds based on pre-designs may also lead to application in micro-tissue engineering. The conclusion implicated that the heterogeneous structures of tissues and organs might be partially derived from the isotropic nature of cell mechanics.

**Figure 7.**
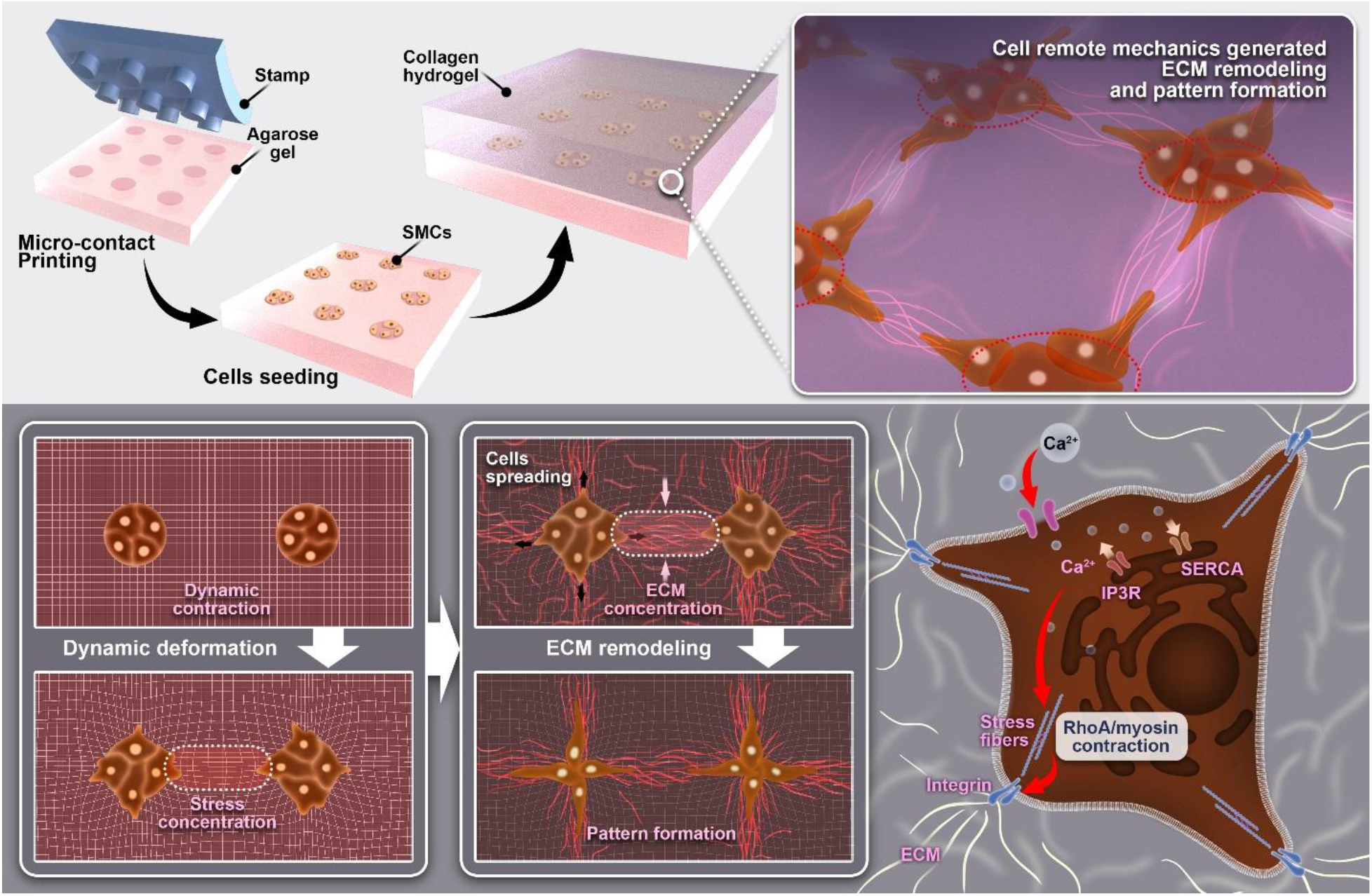
Illustration for remote induction of collagen hydrogel modeling by cell dynamic mechanics. Molecular Dynamics simulations show isotropic stress concentrations and collagen fiber clustering by cell mechanics, along with associated cellular biomechanical mechanism, which implicates the heterogeneous organization of tissues maybe partially originated from the isotropic nature of cell mechanics. (Detailed descriptions seen in the text of the Discussion.)

## Acknowledgements

The work was assisted with help from Dr. Jia Guo, Jingjing Li, Yan Pan, and Lei Liu, and the graphical drawing in Figure 7 by courtesy of Yang Jin (Changzhou University). This project was supported financially by National Natural Science Foundation of China (NSFC 11872129), Natural Science Foundation of Jiangsu Province (BK20181416), Projects of “Jiangsu Specially-appointed Professor” and “Jiangsu six talent peaks (C)” (M.O.); National Natural Science Foundation of China (11902051) (B.B.); National Natural Science Foundation of China (11532003) (L.D.).

## Author contributions

M.O. and L.D. designed the research; B.B. and B.J. designed the computational programs and performed the stimulation work; Y.H. performed the majority of experiments; W.C., H.L., Y.J. and S.Q. helped with experiments and data analysis; L.D. provided the setups of equipment; M.O., Y.H., B.B., L.D. and B.J. prepared the paper.

## Statement for No Conflict of Interest

All authors of this paper declare no conflict of interest in this work.

## References

Abhilash, A.S., B.M. Baker, B. Trappmann, C.S. Chen, and V.B. Shenoy. 2014. Remodeling of fibrous extracellular matrices by contractile cells: predictions from discrete fiber network simulations. Biophysical journal. 107:1829–1840.

Aghvami, M., V.H. Barocas, and E.A. Sander. 2013. Multiscale mechanical simulations of cell compacted collagen gels. Journal of biomechanical engineering. 135:71004.

Alford, A.I., K.M. Kozloff, and K.D. Hankenson. 2015. Extracellular matrix networks in bone remodeling. The international journal of biochemistry & cell biology. 65:20–31.

Alisafaei, F., X. Chen, T. Leahy, P.A. Janmey, and V.B. Shenoy. 2021. Long-range mechanical signaling in biological systems. Soft matter. 17:241–253.

Allen, D.L., and L.A. Leinwand. 2002. Intracellular calcium and myosin isoform transitions. Calcineurin and calcium-calmodulin kinase pathways regulate preferential activation of the IIa myosin heavy chain promoter. The Journal of biological chemistry. 277:45323–45330.

Aoki, K., Y. Kondo, H. Naoki, T. Hiratsuka, R.E. Itoh, and M. Matsuda. 2017. Propagating Wave of ERK Activation Orients Collective Cell Migration. Developmental cell. 43:305–317 e305.

Aper, S.J., A.C. van Spreeuwel, M.C. van Turnhout, A.J. van der Linden, P.A. Pieters, N.L. van der Zon, S.L. de la Rambelje, C.V. Bouten, and M. Merkx. 2014. Colorful protein-based fluorescent probes for collagen imaging. PloS one. 9:e114983.

Baker, B.M., B. Trappmann, W.Y. Wang, M.S. Sakar, I.L. Kim, V.B. Shenoy, J.A. Burdick, and C.S. Chen. 2015. Cell-mediated fibre recruitment drives extracellular matrix mechanosensing in engineered fibrillar microenvironments. Nature materials. 14:1262–1268.

Barnes, C., L. Speroni, K.P. Quinn, M. Montevil, K. Saetzler, G. Bode-Animashaun, G. McKerr, I. Georgakoudi, C.S. Downes, C. Sonnenschein, C.V. Howard, and A.M. Soto. 2014. From single cells to tissues: interactions between the matrix and human breast cells in real time. PloS one. 9:e93325.

Batters, C., D. Brack, H. Ellrich, B. Averbeck, and C. Veigel. 2016. Calcium can mobilize and activate myosin-VI. Proceedings of the National Academy of Sciences of the United States of America. 113:E1162–1169.

Beca, K.I.K., B.M. Girard, T.J. Heppner, G.W. Hennig, G.M. Herrera, M.T. Nelson, and M.A. Vizzard. 2021. The Role of PIEZO1 in Urinary Bladder Function and Dysfunction in a Rodent Model of Cyclophosphamide-Induced Cystitis. Front Pain Res (Lausanne). 2:748385.

Brownfield, D.G., G. Venugopalan, A. Lo, H. Mori, K. Tanner, D.A. Fletcher, and M.J. Bissell. 2013. Patterned collagen fibers orient branching mammary epithelium through distinct signaling modules. Curr Biol. 23:703–709.

Cassereau, L., Y.A. Miroshnikova, G. Ou, J. Lakins, and V.M. Weaver. 2015. A 3D tension bioreactor platform to study the interplay between ECM stiffness and tumor phenotype. J Biotechnol. 193:66–69.

Davidson, C.D., D.K.P. Jayco, W.Y. Wang, A. Shikanov, and B.M. Baker. 2020. Fiber Crimp Confers Matrix Mechanical Nonlinearity, Regulates Endothelial Cell Mechanosensing, and Promotes Microvascular Network Formation. Journal of biomechanical engineering. 142.

Duan, Y., J. Long, J. Chen, X. Jiang, J. Zhu, Y. Jin, F. Lin, J. Zhong, R. Xu, L. Mao, and L. Deng. 2016. Overexpression of soluble ADAM33 promotes a hypercontractile phenotype of the airway smooth muscle cell in rat. Experimental cell research. 349:109–118.

Erdogan, B., M. Ao, L.M. White, A.L. Means, B.M. Brewer, L. Yang, M.K. Washington, C. Shi, O.E. Franco, A.M. Weaver, S.W. Hayward, D. Li, and D.J. Webb. 2017. Cancer-associated fibroblasts promote directional cancer cell migration by aligning fibronectin. J Cell Biol. 216:3799–3816.

Fan, Q., Y. Zheng, X. Wang, R. Xie, Y. Ding, B. Wang, X. Yu, Y. Lu, L. Liu, Y. Li, M. Li, Y. Zhao, Y. Jiao, and F. Ye. 2021. Dynamically Re-Organized Collagen Fiber Bundles Transmit Mechanical Signals and Induce Strongly Correlated Cell Migration and Self-Organization. Angew Chem Int Ed Engl. 60:11858–11867.

Fang, X., K. Ni, J. Guo, Y. Li, Y. Zhou, H. Sheng, B. Bu, M. Luo, M. Ouyang, and L. Deng. 2022. FRET Visualization of Cyclic Stretch-Activated ERK via Calcium Channels Mechanosensation While Not Integrin beta1 in Airway Smooth Muscle Cells. Front Cell Dev Biol. 10:847852.

Guo, C.L., M. Ouyang, J.Y. Yu, J. Maslov, A. Price, and C.Y. Shen. 2012. Long-range mechanical force enables self-assembly of epithelial tubular patterns. Proceedings of the National Academy of Sciences of the United States of America. 109:5576–5582.

Hall, M.S., F. Alisafaei, E. Ban, X. Feng, C.Y. Hui, V.B. Shenoy, and M. Wu. 2016. Fibrous nonlinear elasticity enables positive mechanical feedback between cells and ECMs. Proceedings of the National Academy of Sciences of the United States of America. 113:14043–14048.

Han, W., S. Chen, W. Yuan, Q. Fan, J. Tian, X. Wang, L. Chen, X. Zhang, W. Wei, R. Liu, J. Qu, Y. Jiao, R.H. Austin, and L. Liu. 2016. Oriented collagen fibers direct tumor cell intravasation. Proceedings of the National Academy of Sciences of the United States of America. 113:11208–11213.

Han, Y.L., P. Ronceray, G. Xu, A. Malandrino, R.D. Kamm, M. Lenz, C.P. Broedersz, and M. Guo. 2018. Cell contraction induces long-ranged stress stiffening in the extracellular matrix. Proceedings of the National Academy of Sciences of the United States of America. 115:4075–4080.

Harris, A.K., D. Stopak, and P. Wild. 1981. Fibroblast traction as a mechanism for collagen morphogenesis. Nature. 290:249–251.

He, S., C. Liu, X. Li, S. Ma, B. Huo, and B. Ji. 2015. Dissecting Collective Cell Behavior in Polarization and Alignment on Micropatterned Substrates. Biophysical journal. 109:489–500.

Humphrey, W., A. Dalke, and K. Schulten. 1996. VMD: visual molecular dynamics. J Mol Graph. 14:33–38, 27-38.

Janke, C., and M.M. Magiera. 2020. The tubulin code and its role in controlling microtubule properties and functions. Nature reviews. Molecular cell biology. 21:307–326.

Jung, W.H., N. Yam, C.C. Chen, K. Elawad, B. Hu, and Y. Chen. 2020. Force-dependent extracellular matrix remodeling by early-stage cancer cells alters diffusion and induces carcinoma-associated fibroblasts. Biomaterials. 234:119756.

Kim, T.J., C. Joo, J. Seong, R. Vafabakhsh, E.L. Botvinick, M.W. Berns, A.E. Palmer, N. Wang, T. Ha, E. Jakobsson, J. Sun, and Y. Wang. 2015. Distinct mechanisms regulating mechanical force-induced Ca(2)(+) signals at the plasma membrane and the ER in human MSCs. Elife. 4:e04876.

Kim, T.J., J. Seong, M. Ouyang, J. Sun, S. Lu, J.P. Hong, N. Wang, and Y. Wang. 2009. Substrate rigidity regulates Ca2+ oscillation via RhoA pathway in stem cells. J Cell Physiol. 218:285–293.

Kleinman, H.K., D. Philp, and M.P. Hoffman. 2003. Role of the extracellular matrix in morphogenesis. Current opinion in biotechnology. 14:526–532.

Lee, W., H.A. Leddy, Y. Chen, S.H. Lee, N.A. Zelenski, A.L. McNulty, J. Wu, K.N. Beicker, J. Coles, S. Zauscher, J. Grandl, F. Sachs, F. Guilak, and W.B. Liedtke. 2014. Synergy between Piezo1 and Piezo2 channels confers high-strain mechanosensitivity to articular cartilage. Proceedings of the National Academy of Sciences of the United States of America. 111:E5114–5122.

Li, X., R. Balagam, T.F. He, P.P. Lee, O.A. Igoshin, and H. Levine. 2017. On the mechanism of long-range orientational order of fibroblasts. Proceedings of the National Academy of Sciences of the United States of America. 114:8974–8979.

Liu, L., H. Yu, H. Zhao, Z. Wu, Y. Long, J. Zhang, X. Yan, Z. You, L. Zhou, T. Xia, Y. Shi, B. Xiao, Y. Wang, C. Huang, and Y. Du. 2020. Matrix-transmitted paratensile signaling enables myofibroblast-fibroblast cross talk in fibrosis expansion. Proceedings of the National Academy of Sciences of the United States of America. 117:10832–10838.

Long, Y., Y. Niu, K. Liang, and Y. Du. 2022. Mechanical communication in fibrosis progression. Trends Cell Biol. 32:70–90.

Nitsan, I., S. Drori, Y.E. Lewis, S. Cohen, and S. Tzlil. 2016. Mechanical communication in cardiac cell synchronized beating. Nat Phys. 12:472-+.

Ouyang, M., Z. Qian, B. Bu, Y. Jin, J. Wang, Y. Zhu, L. Liu, Y. Pan, and L. Deng. 2020. Sensing Traction Force on the Matrix Induces Cell-Cell Distant Mechanical Communications for Self-Assembly. ACS biomaterials science & engineering. 6:5833–5848.

Ouyang, M., J.Y. Yu, Y. Chen, L. Deng, and C.L. Guo. 2021. Cell-extracellular matrix interactions in the fluidic phase direct the topology and polarity of self-organized epithelial structures. Cell Prolif. 54:e13014.

Ouyang, M., Y. Zhu, J. Wang, Q. Zhang, Y. Hu, B. Bu, J. Guo, and L. Deng. 2022. Mechanical communication-associated cell directional migration and branching connections mediated by calcium channels, integrin beta1, and N-cadherin. Front Cell Dev Biol. 10:942058.

Pakshir, P., M. Alizadehgiashi, B. Wong, N.M. Coelho, X. Chen, Z. Gong, V.B. Shenoy, C.A. McCulloch, and B. Hinz. 2019. Dynamic fibroblast contractions attract remote macrophages in fibrillar collagen matrix. Nature communications. 10:1850.

Pankova, D., Y. Chen, M. Terajima, M.J. Schliekelman, B.N. Baird, M. Fahrenholtz, L. Sun, B.J. Gill, T.J. Vadakkan, M.P. Kim, Y.H. Ahn, J.D. Roybal, X. Liu, E.R. Parra Cuentas, J. Rodriguez, Wistuba, II, C.J. Creighton, D.L. Gibbons, J.M. Hicks, M.E. Dickinson, J.L. West, K.J. Grande-Allen, S.M. Hanash, M. Yamauchi, and J.M. Kurie. 2016. Cancer-Associated Fibroblasts Induce a Collagen Cross-link Switch in Tumor Stroma. Mol Cancer Res. 14:287–295.

Reinhart-King, C.A., M. Dembo, and D.A. Hammer. 2008. Cell-cell mechanical communication through compliant substrates. Biophysical journal. 95:6044–6051.

Sapir, L., and S. Tzlil. 2017. Talking over the extracellular matrix: How do cells communicate mechanically? Semin Cell Dev Biol. 71:99–105.

Sawhney, R.K., and J. Howard. 2002. Slow local movements of collagen fibers by fibroblasts drive the rapid global self-organization of collagen gels. J Cell Biol. 157:1083–1091.

Shellard, A., and R. Mayor. 2021. Collective durotaxis along a self-generated stiffness gradient in vivo. Nature. 600:690–694.

Shellard, A., A. Szabo, X. Trepat, and R. Mayor. 2018. Supracellular contraction at the rear of neural crest cell groups drives collective chemotaxis. Science. 362:339–343.

Shi, Q., R.P. Ghosh, H. Engelke, C.H. Rycroft, L. Cassereau, J.A. Sethian, V.M. Weaver, and J.T. Liphardt. 2014. Rapid disorganization of mechanically interacting systems of mammary acini. Proceedings of the National Academy of Sciences of the United States of America. 111:658–663.

Stopak, D., and A.K. Harris. 1982. Connective tissue morphogenesis by fibroblast traction. I. Tissue culture observations. Dev Biol. 90:383–398.

Stopak, D., N.K. Wessells, and A.K. Harris. 1985. Morphogenetic rearrangement of injected collagen in developing chicken limb buds. Proceedings of the National Academy of Sciences of the United States of America. 82:2804–2808.

Sunyer, R., V. Conte, J. Escribano, A. Elosegui-Artola, A. Labernadie, L. Valon, D. Navajas, J.M. Garcia-Aznar, J.J. Munoz, P. Roca-Cusachs, and X. Trepat. 2016. Collective cell durotaxis emerges from long-range intercellular force transmission. Science. 353:1157–1161.

Tanner, K., H. Mori, R. Mroue, A. Bruni-Cardoso, and M.J. Bissell. 2012. Coherent angular motion in the establishment of multicellular architecture of glandular tissues. Proceedings of the National Academy of Sciences of the United States of America. 109:1973–1978.

Theocharis, A.D., D. Manou, and N.K. Karamanos. 2019. The extracellular matrix as a multitasking player in disease. FEBS J. 286:2830–2869.

Ushiki, T. 2002. Collagen fibers, reticular fibers and elastic fibers. A comprehensive understanding from a morphological viewpoint. Arch Histol Cytol. 65:109–126.

Vicente-Manzanares, M., X. Ma, R.S. Adelstein, and A.R. Horwitz. 2009. Non-muscle myosin II takes centre stage in cell adhesion and migration. Nature reviews. Molecular cell biology. 10:778–790.

Wang, H., A.S. Abhilash, C.S. Chen, R.G. Wells, and V.B. Shenoy. 2014. Long-range force transmission in fibrous matrices enabled by tension-driven alignment of fibers. Biophysical journal. 107:2592–2603.

Wang, J., J. Guo, B. Che, M. Ouyang, and L. Deng. 2020. Cell motion-coordinated fibrillar assembly of soluble collagen I to promote MDCK cell branching formation. Biochemical and biophysical research communications. 524:317–324.

Wang, J., S. Zhao, L. Luo, Y. Liu, E. Li, Z. Zhu, and Z. Zhao. 2019. Shengjing Capsule Improves Spermatogenesis through Upregulating Integrin alpha6/beta1 in the NOA Rats. Evid Based Complement Alternat Med. 2019:8494567.

Wennberg, C.L., T. Murtola, S. Pall, M.J. Abraham, B. Hess, and E. Lindahl. 2015. Direct-Space Corrections Enable Fast and Accurate Lorentz-Berthelot Combination Rule Lennard-Jones Lattice Summation. J Chem Theory Comput. 11:5737–5746.

Winer, J.P., S. Oake, and P.A. Janmey. 2009. Non-linear elasticity of extracellular matrices enables contractile cells to communicate local position and orientation. PloS one. 4:e6382.

Wu, Y., Z. Jiang, X. Zan, Y. Lin, and Q. Wang. 2017. Shear flow induced long-range ordering of rod-like viral nanoparticles within hydrogel. Colloids Surf B Biointerfaces. 158:620–626.

Yao, M., A. Tijore, D. Cheng, J.V. Li, A. Hariharan, B. Martinac, G. Tran Van Nhieu, C.D. Cox, and M. Sheetz. 2022. Force- and cell state-dependent recruitment of Piezo1 drives focal adhesion dynamics and calcium entry. Science advances. 8:eabo1461.

Zarkoob, H., S. Chinnathambi, J.C. Selby, and E.A. Sander. 2018. Substrate deformations induce directed keratinocyte migration. J R Soc Interface. 15.

