## Supporting information for "Cell Dynamic Mechanics Regulates Large-spatial Isotropic Matrix Modeling with Computational Simulations"

The supplementary information contains 1 movie and 10 figures.

### Supplementary movie and figures

**Movie 1:** The dynamic process by Molecular Dynamics simulation shows the fiber emergences in connecting the cell clusters in random-style array in corresponding to the data of Figure 2I.

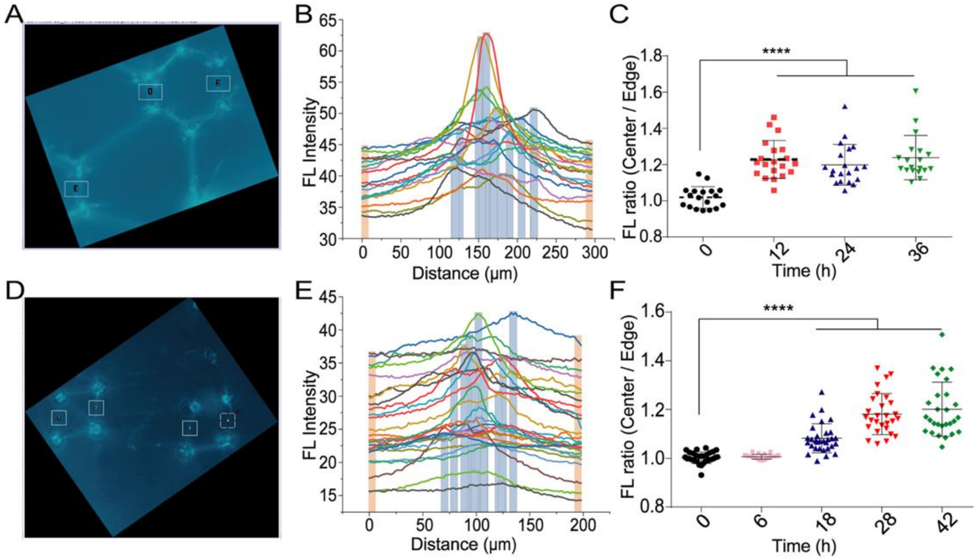


**Figure S1. The procedural demo for COL fluorescence quantifications.** The processing details with text were described in the Methods. (**A-C**) The sample of single hexagon for COL fluorescence quantifications. (A) the COL fluorescence image along with selected regions (ROI); (B) the fluorescence distributions of individual ROIs; (C) fluorescence-intensity (FL) ratio quantifications (mean ± S.E.M.) at the different time points. (**D-F**) The sample of single parallelogram for the fluorescence quantifications.

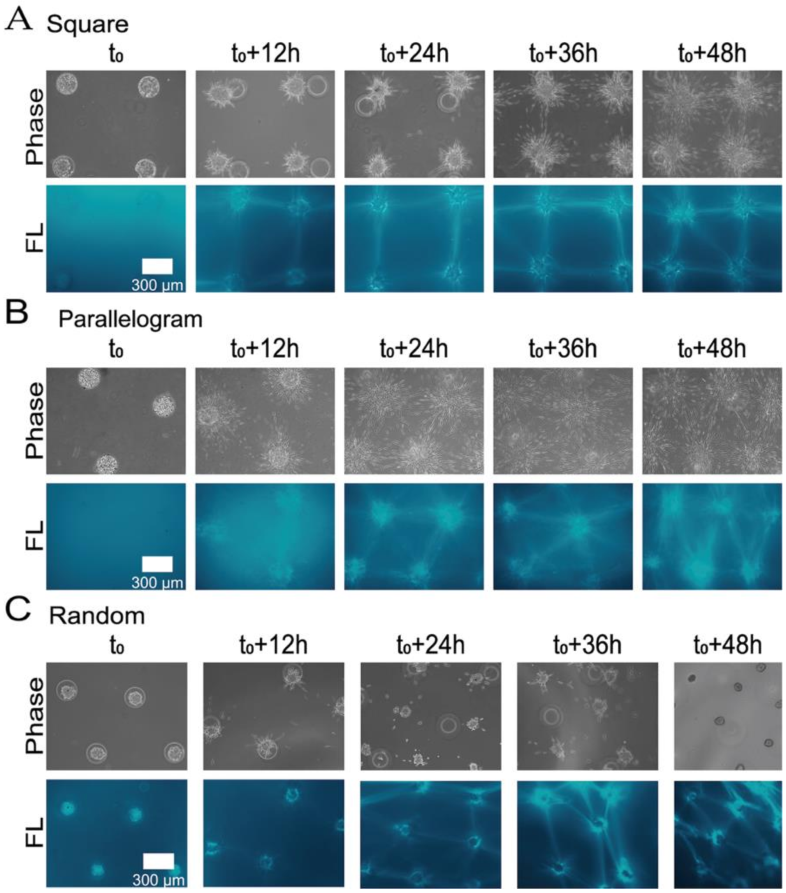


**Figure S2. The COL fiber inductions along the culture times in different geometrical arrays.** The images were taken with x10 objective, in corresponding to the images taken with x5 objective in Figure 2. **(A-C)** The representative bright-field and COL fluorescence images along the culture time in large-spatial square (A), parallelogram (B), and random-style (C) arrays.

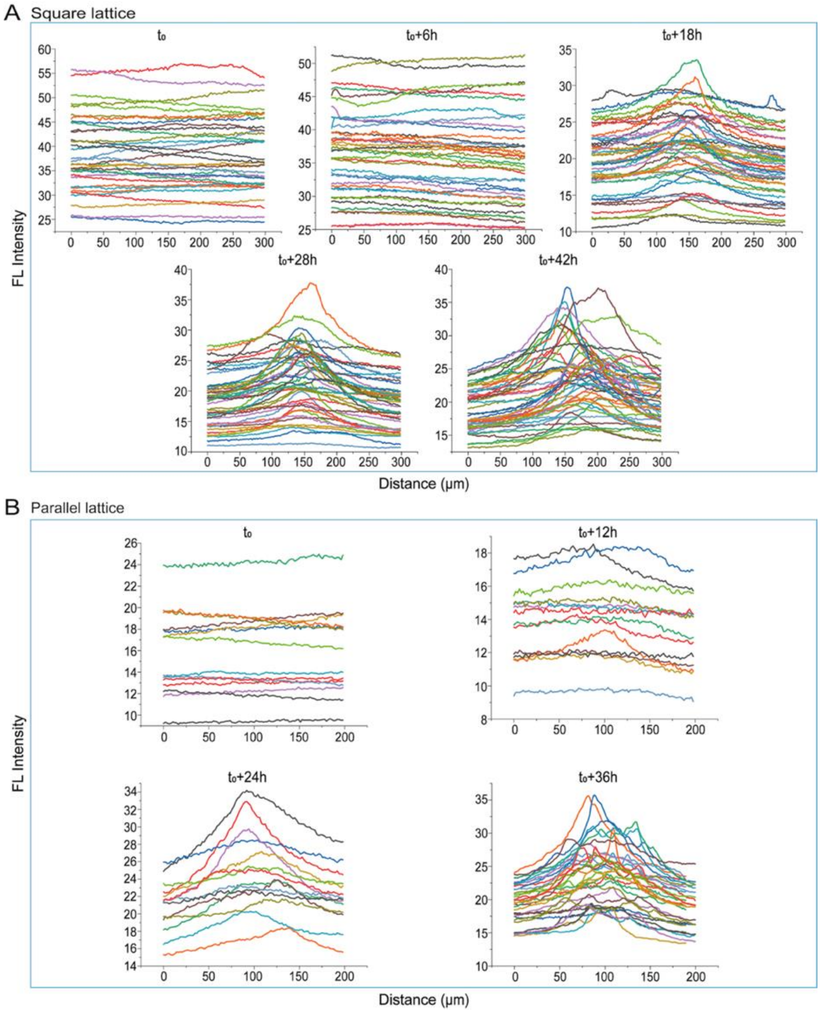


**Figure S3. COL fluorescence distribution curves along the individual selected regions.** As representative demonstrations, the quantifications of fluorescence (FL) curves showed the gradual emerging COL fibers at the different culture time. **(A, B)** The curves of COL fluorescence distributions at the large-spatial square arrays (A), and parallelogram arrays (B).

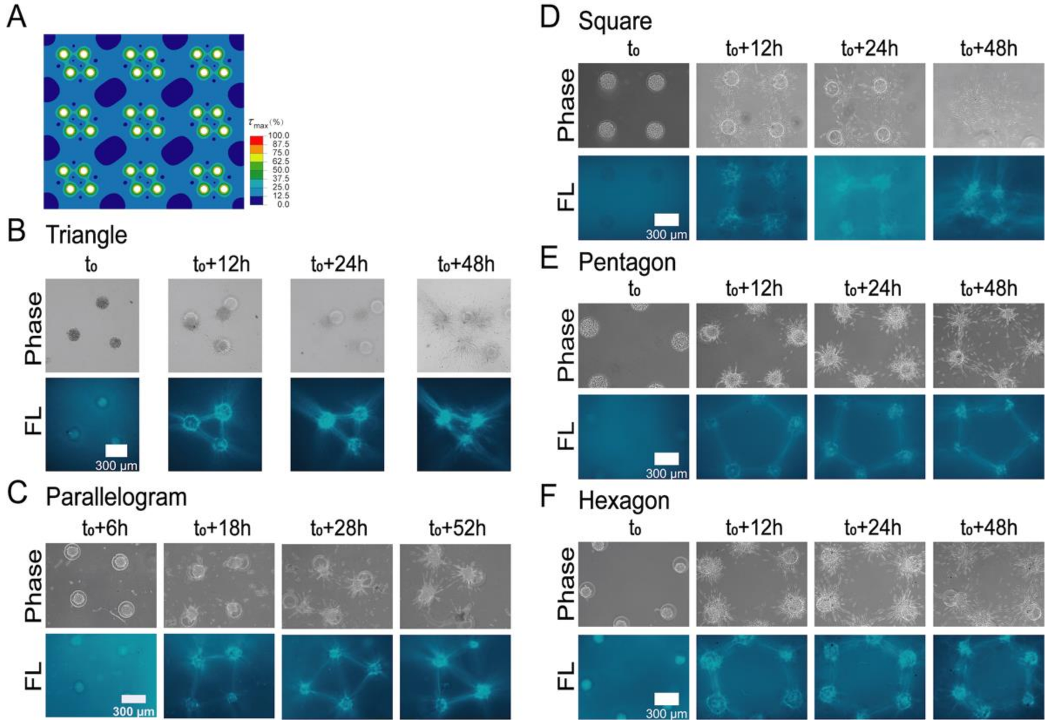


**Figure S4. COL fiber inductions in single polygons with diverse geometries.** The images were taken with x10 objective, in corresponding to Figure 3. **(A)** The maximum stress distribution in single parallelogram by FEM stimulation. **(B-F)** The representative bright-field and COL fluorescence images from single triangles to hexagons.

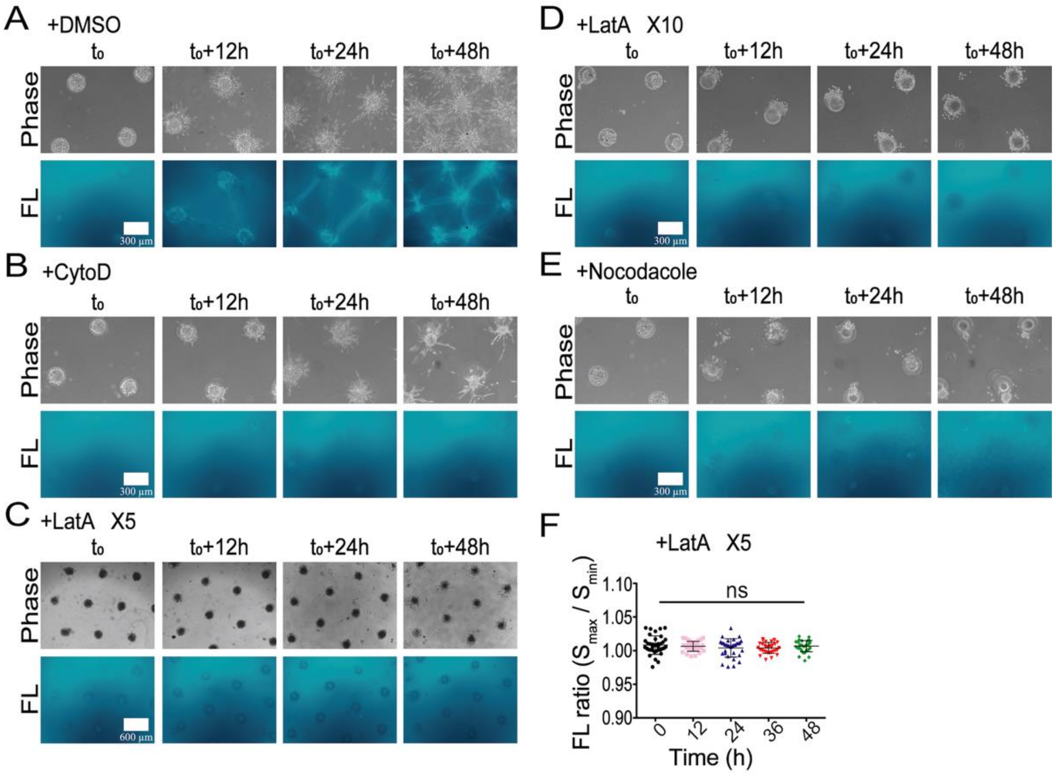


**Figure S5. The importance of cellular cytoskeleton integrity in remote COL fiber inductions.** The images were taken with x10 or x5 objective, in corresponding to Figure 4(A-D). **(A-F)** The representative images at the different culture time under control condition (DMSO) or treated with CytoD, LatA or Nocodazole (A-D), and fluorescence quantification for LatA treatment (F).

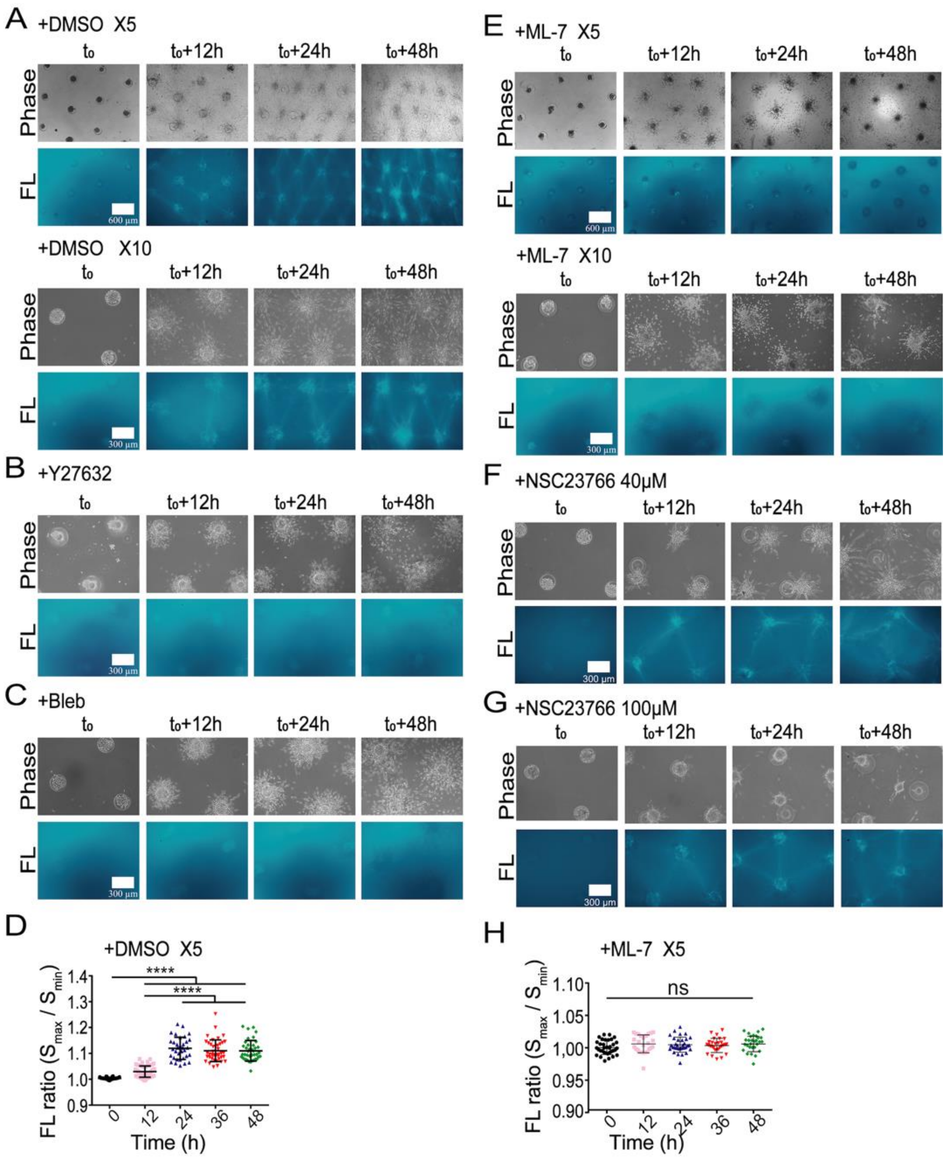


**Figure S6. The importance of actomyosin contraction in remote COL fiber inductions.** The images were taken with x10 or x5 objective, in corresponding to Figure 4(E-H). **(A-H)** The representative images at the different culture time under control condition (DMSO) or treated with Y27632, Blebbistanin, ML-7, or NSC23766 at 40 or 100 μM, and fluorescence quantification for DMSO (D) or ML-7 (H) treatment.

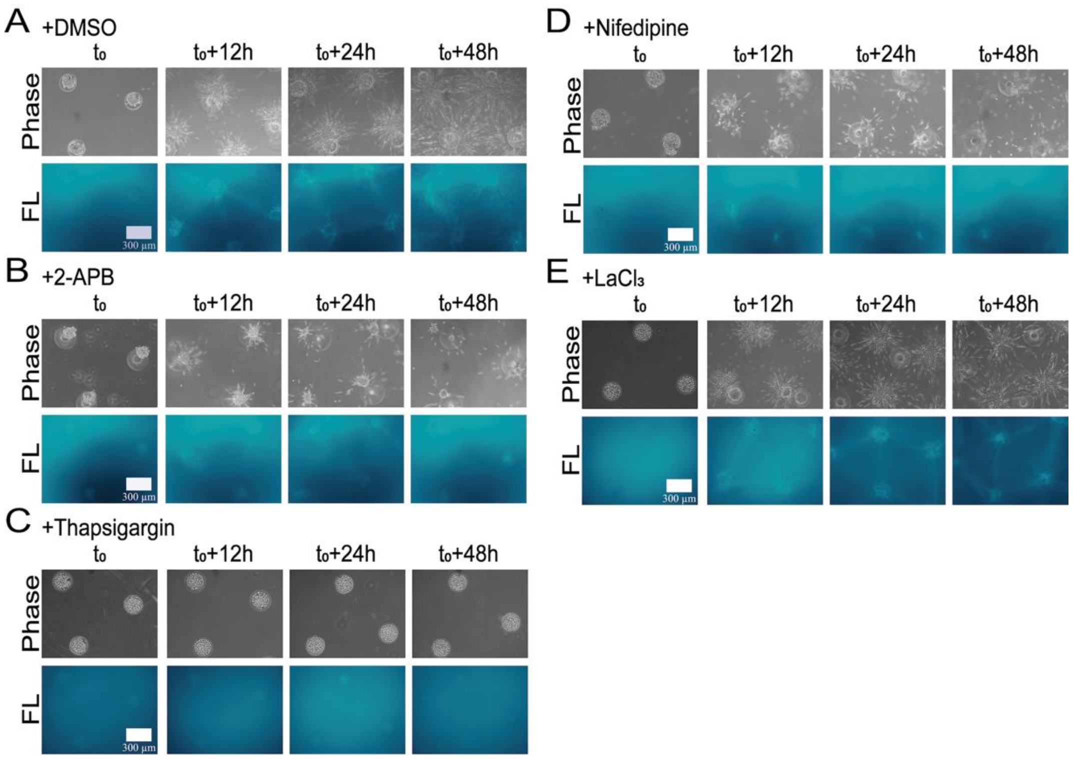


**Figure S7. Measurements of COL fiber inductions under inhibitions of variable calcium channels.** The images were taken with x10 objective, in corresponding to Figure 5. **(A-E)** The representative bright-field and COL fluorescence images under control (DMSO) or treated with 2-APB, Thapsigargin, Nifedipine, or $LaCl_3$.

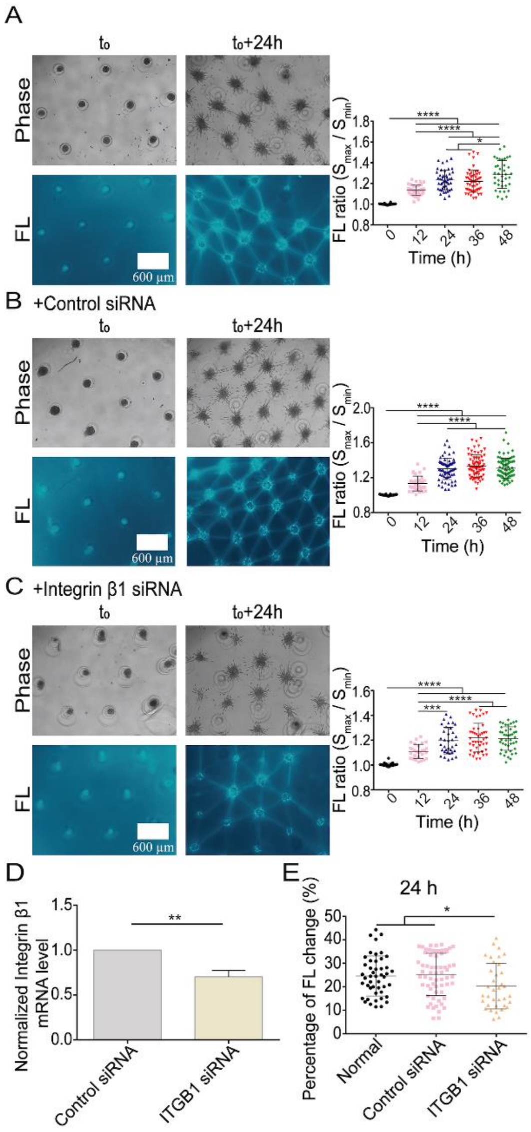


**Figure S8. The remote COL fiber inductions under transfection with ITGB1 siRNA.** ASM cells were transfected with positive or negative siRNA (50 nM) for integrin β1 (ITGB1) or Piezo1. COL fiber inductions were recorded and quantified in geometrical parallelogram-arrayed clusters. **(A-C)** Fluorescent COL fibers and quantifications under the condition of cell clusters without transfection (**A**), or with negative siRNA (**B**) or ITGB1 siRNA (**C**) transfection. **(D, E)** The mRNA expression levels (**D**) and comparison of percentage changes in fiber growth (**E**) with ITGB1 siRNA transfection.

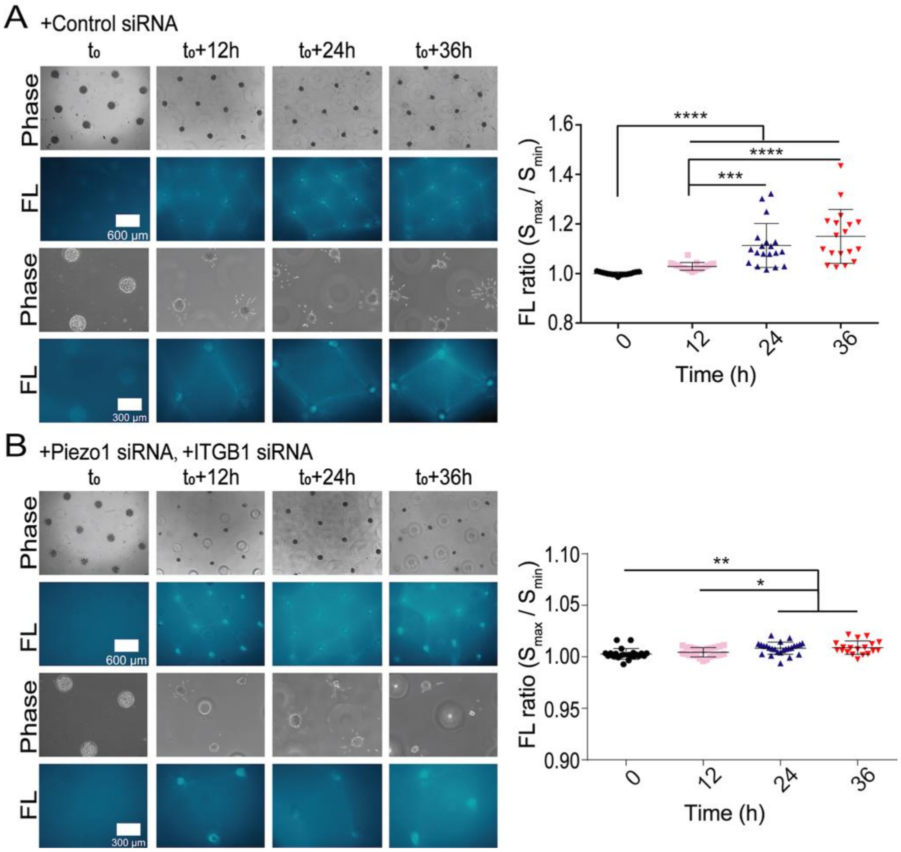


**Figure S9. The role of double down-regulations of integrin β1 and Piezo1 by siRNA transfection in COL fiber inductions.** The images were taken with x10 objective, in corresponding to images in Figure 6(A, B). **(A, B)** The representative bright-field and COL fluorescence images transfected with negative siRNA or double ITGB1 and Piezo1 siRNA.

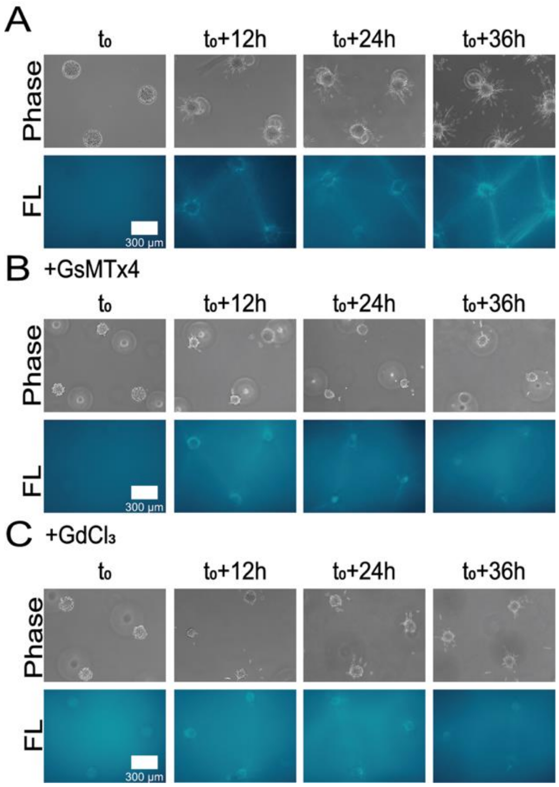


**Figure S10. The role of Piezo inhibition in COL fiber inductions.** The images were taken with x10 objective, in corresponding to images in Figure 6(D-F). **(A-C)** The representative bright-field and COL fluorescence images under control condition, or treated with GsMTx4 or $GdCl_3$.
